# Resolution Enhancement with a Task-Assisted GAN to Guide Optical Nanoscopy Image Analysis and Acquisition

**DOI:** 10.1101/2021.07.19.452964

**Authors:** Catherine Bouchard, Theresa Wiesner, Andréanne Deschênes, Anthony Bilodeau, Benoît Turcotte, Christian Gagné, Flavie Lavoie-Cardinal

## Abstract

We introduce a deep learning model that predicts super-resolved versions of diffraction-limited microscopy images. Our model, named Task- Assisted Generative Adversarial Network (TA-GAN), incorporates an auxiliary task (e.g. segmentation, localization) closely related to the observed biological nanostructures characterization. We evaluate how TA-GAN improves generative accuracy over unassisted methods using images acquired with different modalities such as confocal, brightfield (diffraction-limited), super-resolved stimulated emission depletion, and structured illumination microscopy. The generated synthetic resolution enhanced images show an accurate distribution of the F-actin nanostructures, replicate the nanoscale synaptic cluster morphology, allow to identify dividing S. aureus bacterial cell boundaries, and localize nanodomains in simulated images of dendritic spines. We expand the applicability of the TA-GAN to different modalities, auxiliary tasks, and online imaging assistance. Incorporated directly into the acquisition pipeline of the microscope, the TA-GAN informs the user on the nanometric content of the field of view without requiring the acquisition of a super-resolved image. This information is used to optimize the acquisition sequence, and reduce light exposure. The TA-GAN also enables the creation of domain-adapted labeled datasets requiring minimal manual annotation, and assists microscopy users by taking online decisions regarding the choice of imaging modality and regions of interest.

## Introduction

The development of super-resolution optical microscopy (optical nanoscopy) techniques to study the nanoscale organisation of biological structures has transformed our understanding of cellular and molecular processes [1]. Such techniques, including STimulated Emission Depletion (STED) microscopy [2], are compatible with live-cell imaging, enabling the monitoring of sub-cellular dynamics with unprecedented spatio-temporal precision. In the design of optical nanoscopy experiments, multiple and often conflicting objectives (e.g. spatial resolution, acquisition speed, light exposure, and signal-to-noise ratio) must be considered [3, 4]. Machine learning-assisted microscopy approaches have been proposed to improve the acquisition processes, mostly by limiting light exposure [3, 5, 6]. In parallel, several supervised [7–11] and weakly supervised [12–14] deep learning approaches have been developed for high-throughput analysis of microscopy images. Deep learning-based super-resolution [5, 15–18] and domain adaptation [19] approaches have also been proposed recently for optical microscopy, but concerns and skepticism arise regarding their applicability to characterize biological structures at the nanoscale [20–22].

Optical nanoscopy techniques exploit the ability to modulate the emission properties of fluorescent molecules to overcome the diffraction limit of light microscopy [23]. In this context, it is challenging to rely on algorithmic methods to generate images of sub-diffraction structures that are not optically resolved in the original image [20]. Methods that are optimized for generating images that *appear* to belong to the target higher-resolution domain do not specifically guarantee that the biological features of interest are accurately generated [22]. Yet, the possibility to super-resolve microscopy images post-acquisition would favorably alleviate some of the compromises between the acquisition parameters in optical nanoscopy [16, 24].

Among the methods developed for algorithmic super-resolution, conditional Generative Adversarial Networks (cGAN) [25] generate data instances based on a different input value, capturing some of its features to guide the creation of a new instance that fits the target domain. However, realism of the synthetic images does not ensure that the images are usable for further field-specific analysis, which is limiting their use in optical microscopy. The primary goal for generating super-resolved microscopy images is to produce reliable nanoscale information on the biological structures of interest. Optimizing a network using auxiliary tasks, or multi-task learning, can guide the generator to resolve content that matters for the current context [26]. Various applications of cGANs for image-to-image translation use auxiliary tasks such as semantic segmentation [27, 28], attributes segmentation [29], or foreground segmentation [30], to provide spatial guidance to the generator. We adapt this idea in the context of microscopy, where structure-specific annotations can direct the attention to subtle features that are only recognizable by trained experts.

We propose to guide the image generation process using an auxiliary task that is closely related to the biological question at hand. This approach improves the applicability of algorithmic super-resolution and ensures that the generated features in synthetic images are consistent with the observed biological structures in real nanoscopy images. Microscopy image analysis tasks that are already routinely solved with deep learning [10] (e.g. segmentation, detection, and classification) can guide a cGAN to preserve the biological features of interest in the generated synthetic images. We introduce a Task-Assisted GAN (TA-GAN) for resolution enhanced microscopy image generation. The TAGAN relies on an auxiliary task associated with structures that are unresolved by the input low-resolution modalities (e.g. confocal or brightfield microscopy) but are easily distinguishable in the targeted super-resolution modalities (e.g. STED or Structured Illumination Microscopy (SIM)). We expand the applicability of the method with a variation called TA-CycleGAN, based on the CycleGAN model [31], applicable to unpaired datasets. Here, the TA-CycleGAN is applied to domain adaptation for STED microscopy of fixed and living neurons. Our results demonstrate that the TA-GAN and TA-CycleGAN models improve the synthetic representation of biological nanostructures in comparison to other algorithmic super-resolution approaches. Specifically, our method is useful to 1) guide the quantitative analysis of nanostructures, 2) generate synthetic datasets of different modalities for data augmentation or to reduce the annotation burden, and 3) predict regions of interest for machine learning-assisted live-cell STED imaging.

## Results

### TA-GAN: Task-assisted super-resolution image generation

Deep learning methods designed for synthetic microscopy image generation have been shown to be effective for deblurring and denoising confocal images [15, 16, 18]. To increase the accuracy of resolution enhancement approaches applied to the generation of complex nanoassemblies, we consider the combination of a cGAN with an additional convolutional neural network, the task network (Figure 1a), targeting an image analysis task relevant to the biological structures of interest. Three individual networks form the TAGAN model: 1) the generator, 2) the discriminator, and 3) the task network (Figure 1a). The chosen auxiliary task should be achievable using the high-resolution modality only, ensuring that it is informative about content that is not resolved in the low-resolution input modality. The error between the task network predictions and the ground truth annotations is back-propagated to the generator to optimize its parameters (Methods). The TA-GAN is trained using pairs of low-resolution (confocal or brightfield) and super-resolution (STED or SIM) images.

**Fig. 1:**
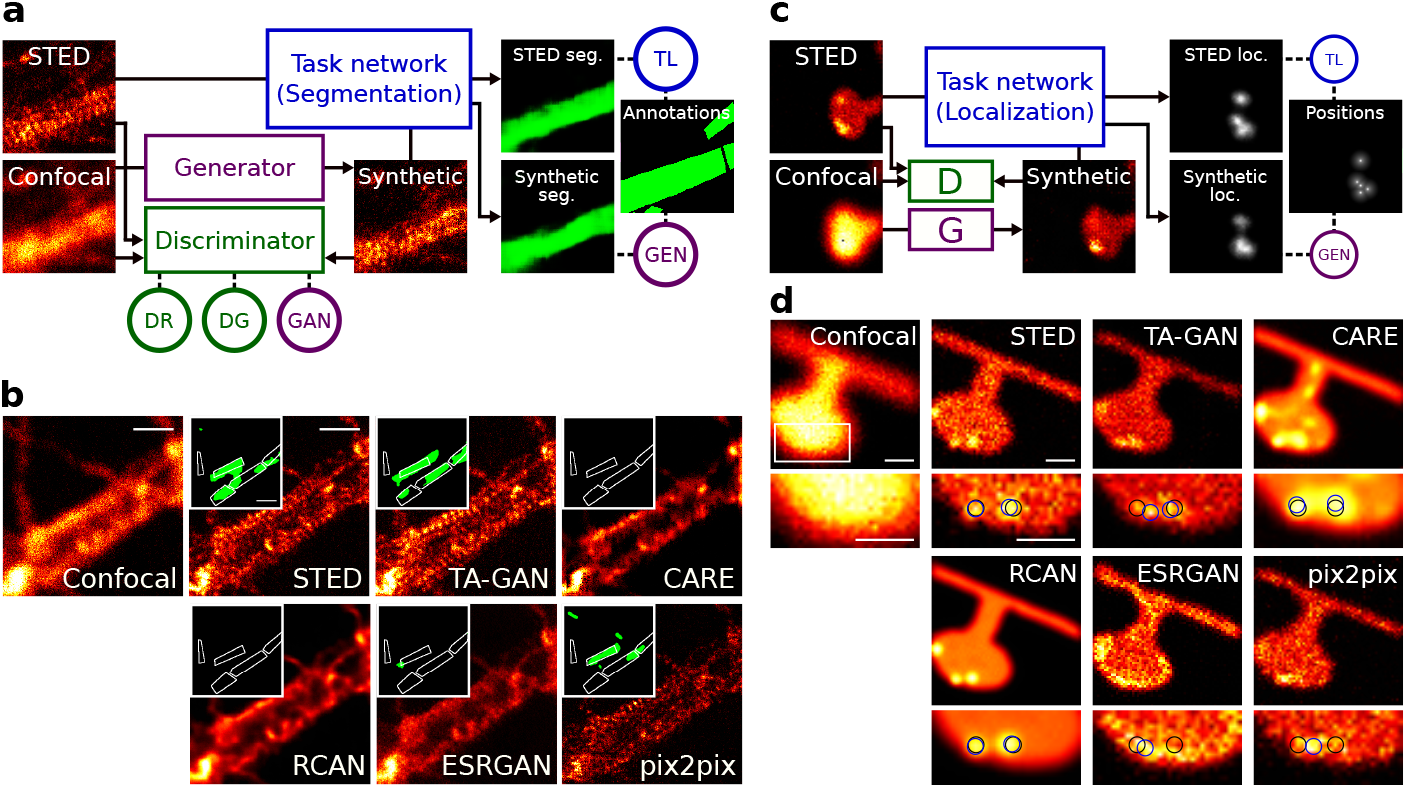
The TA-GAN method. **a,** Architecture of the TA-GAN*_Ax._*. The losses (circles) are backpropagated to the networks of the same color: the generator (violet), the discriminator (green), and the task network (blue). (DG: Discriminator loss for generated images, GEN: Generation loss, GAN: GAN loss, DR: Discriminator loss for real images, TL: Task loss.). The TA-GAN*_Ax._* is applied to the *Axonal F-actin dataset* using the segmentation of F-actin rings as an auxiliary task to optimize the generator. **b,** Comparison of the TA-GAN*_Ax._* and algorithmic super-resolution baselines on the *Axonal F-actin dataset*. The confocal image is the low-resolution input and the STED image is the aimed ground truth. Inset: segmentation of the axonal F-actin rings (green) predicted by the U-Net*_Fixed−ax._* with the bounding boxes (white line) corresponding to the manual expert annotations [14]. Scale bars: 1 µm. **c,** The TAGAN*_Nano._* is trained on the *Simulated nanodomain dataset* using the localization of nanodomains as the auxiliary task. **d,** Comparison of the TA-GAN*_Nano._* with the baselines for nanodomain localization. The black circles represent the position of the nanodomains on the ground truth datamap and the blue circles represent the nanodomains identified by an expert on images from the test set (Methods). The intensity scale is normalized for each image by its respective minimum and maximum values. Scale bars: 250 nm.

The first TA-GAN model, TA-GAN*_Ax._*, is trained on the *Axonal F-actin dataset* to generate STED images of the axonal F-actin lattice from confocal images (Figure 1b). The auxiliary task identified to train the TA-GAN*_Ax._* is the segmentation of the axonal F-actin rings, which cannot be resolved with confocal microscopy [32] (Figure 1a). The segmentation network output is used to compute the generation loss and to evaluate the generation performance at test time. The image super-resolution baselines CARE [16], RCAN [15], ESRGAN [33, 34], and pix2pix [35], are trained on the *Axonal F-actin dataset* and applied to the generation of a synthetic resolution enhanced image from an input confocal image (Figure 1b, first row). We additionally evaluate the performance of image denoising baselines DnCNN [36, 37] and Noise2Noise [37, 38] on the confocal to STED image translation task (Supplementary Figure 1). Comparison between the results of the TA-GAN*_Ax._* with the baselines reveals that the pixel-wise mean square error (MSE), structural similarity index (SSIM), and peak signal-to-noise ratio (PSNR) between generated and ground truth (GT) STED images are either improved or similar using the TA-GAN*_Ax._* (Extended data fig. 1, Supplementary Figure 2, and Supplementary Figure 3). To evaluate the accuracy of each baseline in the generation of the nanostructure of interest, we evaluate the ability of an independent deep learning model trained on real STED images only [14], which we refer to as U-Net*_F ixed−ax._*, to segment the F-actin rings in the synthetic images. The TAGAN*_Ax._* model uses the segmentation loss to optimize the generator’s weights, which forces the generated F-actin nanostructures to be realistic enough to be recognized by the task network during training, and U-Net*_F ixed−ax._* during testing. The U-Net*_F ixed−ax._* is applied to the synthetic and real STED images, and the similarity between the resulting pairs of segmentation maps is computed using the Dice coefficient (DC) and Intersection over Union (IOU) metrics. The improvement in similarity is significant for TA-GAN*_Ax._* compared to all baselines (Extended data fig. 1).

We created a dataset of nanodomains in simulated shapes of dendritic spines using the *pySTED* simulation platform [39] to characterize the conditions where the TA-GAN outperforms the baselines in a controlled environment. The task used to train the TA-GAN*_Nano._* is the localization of the centers of the simulated nanodomains (Figure 1c). We compare the generated images with the ground truth datamaps for two analysis tasks : 1) the localization of two nanodomains that are spaced by less than 100 nm, which is too close to be resolved with a standard deconvolution approach (Richardson Lucy [40]), and 2) the counting of nanodomains (2 to 6) separated by variable distances. The localization of the nanodomains can be performed using the TA-GAN synthetic images with similar accuracy to the one obtained using the simulated STED images from the pySTED platform (Supplementary Figure 5a). For the counting task, the images generated by the TA-GAN, RCAN, and pix2pix, allow to count up to six nanodomains that cannot be resolved in the simulated confocal images within a simulated spine (Supplementary Figure 5b). Similarly to the results obtained on the *Axonal F-actin dataset*, TA-GAN and pix2pix are the two algorithmic super-resolution approaches that generate synthetic images with the highest similarity to the target domain for the *Simulated nanodomain dataset*, preserving image features such as the signal-to-noise ratio, background level, and spatial resolution(Figure 1d, and Supplementary Figure

The TA-GAN model requires the definition of a task that steers the training of the generator toward the accurate extraction of sub-resolution information. The addition of this task is what differentiates TA-GAN from baselines like pix2pix. We therefore evaluate how the choice of task impacts the performance using two different datasets. For the *Synaptic Proteins dataset* [41], we evaluate the approach using a localization (Figure 2a) and a segmentation task (Supplementary Figure 6). The annotations are automatically generated using the pySODA analysis strategy [41]. For the localization task we use the weighted centroids of the clusters, while for the segmentation task the masks are generated with wavelet segmentation [42]. We show that both tasks can be used to guide the synthetic image generation (Figure 2b,c), but that the localization task allows to generate synaptic protein clusters with morphological features that are more similar to the one observed in the real images (Supplementary Figure 7 and Supplementary Figure 8).

**Fig. 2:**
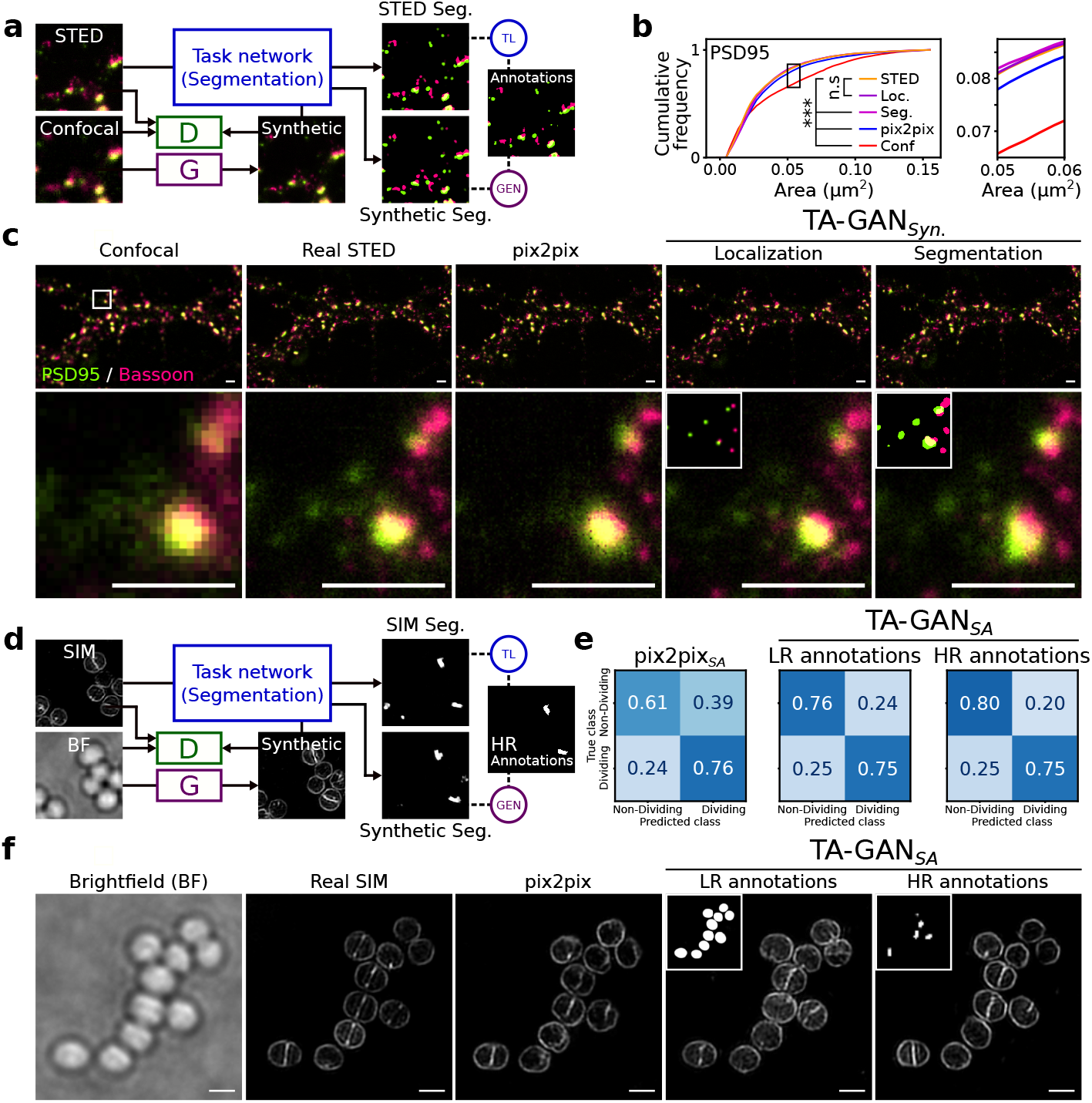
Dataset specific tasks drive reliable resolution enhancement with the TA-GAN approach. **a,** Two TA-GAN models designed for the *Synaptic protein dataset* are trained using one of two auxiliary tasks: the segmentation of the protein clusters (shown) or the localization of the weighted centroids (Supplementary Figure 6). **b,** Comparison between the different approaches for the characterization of synaptic cluster morphological features. Shown is the cumulative distribution of the cluster area for PSD95 (see Supplementary Figure 7 for more analysed features). Statistical analysis: two-sample Kolmogorov–Smirnov test. **c,** Generation of synthetic two-color images of PSD95 and Bassoon on the test set of the *Synaptic protein dataset* using the non-task-assisted baseline (pix2pix), the TA-GAN*_Syn._* with the localization task, and the TA-GAN*_Syn._* with the segmentation task. Insets: localization and segmentation annotations used to train the two TA-GAN*_Syn._* models. Scale bars: 1 µm. Each crop is normalized to the 98^th^ percentile of its pixel values for better visualization of dim clusters. **d,** The TA-GAN*_SA._* models designed for the *S. aureus dataset* are trained using a segmentation task with annotations requiring only the LR brightfield image or annotations requiring the HR SIM image. **e,** Confusion matrices for the classification of dividing and non-dividing cells on the test set of the *S. aureus dataset* (N=410 cells in 5 images). The TA-GAN*_SA._* trained with HR annotations achieves better performance in generating the boundaries between dividing bacterial cell, a morphological feature visible only with SIM microscopy, compared to pix2pix and the TA-GAN*_SA._* trained with LR annotations. **f,** Images obtained with pix2pix and the TA-GAN*_SA._* trained with LR and HR annotations. Insets: LR and HR annotations used to train the two TA-GAN*_S_A.* models. Scale bars: 1 µm.

We evaluate how the precision of the labels used for the task impacts the generation accuracy using the publicly available dataset of *S. aureus* cells from DeepBacs [43, 44]. S. aureus bacteria are very small (around 1 µm diameter), and monitoring their morphology changes and cell division processes requires sub-diffraction resolution [45]. The TA-GAN*_SA._* is trained for brightfield to SIM resolution enhancement using a classification task based either on: 1) low-resolution (LR) annotations generated from the brightfield modality (Supplementary Figure 6) or 2) high-resolution (HR) annotations of dividing cell boundaries obtained from the SIM modality (Figure 2d). We evaluate how the images generated with the algorithmic super-resolution approaches can be used for the classification of dividing and non-dividing bacterial cells, a task that is not achievable using only brightfield microscopy images (Supplementary Figure 9). Training the TA-GAN*_SA._* model using the HR annotations leads to an improved classification performance combined with improved realism of the synthetic images (Figure 2e,f).

### TA-CycleGAN: Domain adaptation on unpaired datasets

For many microscopy modalities, paired *and labeled* training datasets are not directly available, or would require a high annotation burden from highly qualified experts. Based on the results obtained using confocal and STED image pairs on fixed neurons, we wanted to expand the applicability of the TA-GAN to unpaired datasets – here, images of fixed and living cells. We first validate that the TA-GAN cabe applied to the *Dendritic F-actin dataset* [14] using the semantic segmentation of F-actin rings and fibers in dendrites of fixed neurons (Figure 3a). The trained TA-GAN*_Dend._* generates synthetic nanostructures that are successfully segmented by the U-Net*_F ixed−dend._* which recognizes dendritic F-actin rings and fibers in real STED images [14] (Figure 3b). Similar to results previously obtained from real STED images [14], the segmentation of the synthetic images with U-Net*_F ixed_* shows that the area of the F-actin rings significantly decreases as the neuronal activity increases, while the opposite is observed for F-actin fibers (Supplementary Figure 10). Using our task-assisted strategy, we next trained a CycleGAN [35] model, as it was precisely developed for image domain translation on unpaired datasets. The TA-CycleGAN can be applied to the translation between two microscopy modalities or experimental conditions in which the same biological structure can be observed (here the F-actin cytoskeleton in cultured live and fixed neurons) without the need for paired images. To this aim we generated the *Live F-actin dataset* consisting of confocal and STED images of F-actin nanostructures in living neurons using the far-red fluorogenic dye SiR-Actin [46].

**Fig. 3:**
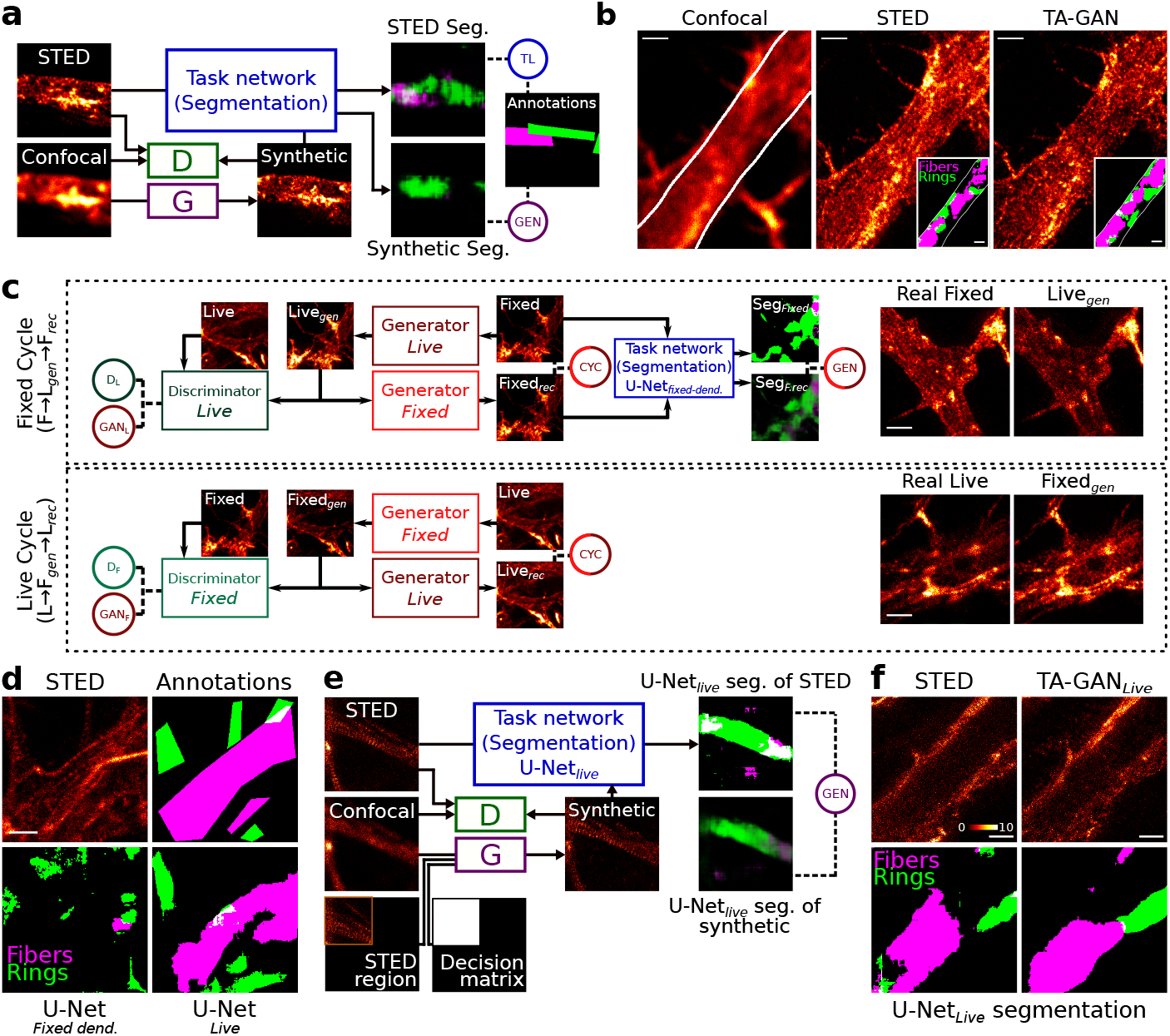
Domain adaptation. **a,** The semantic segmentation of F-actin rings (green) and fibers (magenta) is used as the auxiliary task to train the TA-GAN*_Dend._*. **b,** Confocal, real STED, and TA-GAN*_Dend._* synthetic images. Insets show the regions identified as rings and fibers by the U-Net*_Fixed−dend._* trained on real STED images [14]. **c,** The same semantic segmentation task is used to train the TACycleGAN. The reference to compute the TL is the segmentation of real fixed-cell STED images by U-Net*_Fixed−dend._*. The *Fixed* cycle (top) uses U-Net*_Fixed−dend._* to encourage semantic consistency between the input fixed-cell image and the end-of-cycle reconstructed image. The *Live* cycle (bottom) does not use a task network, enabling the use of non-annotated images from the *Live F-actin dataset*. Once trained, the TACycle-GAN can generate domain adapted datasets (right). **d,** Segmentation of F-actin nanostructures on live-cell STED images. The nanostructures on the live-cell STED images (top-left) are not properly segmented by the U-Net*_Fixed−Dend._* (bottom-left). The U-Net*_Live_* is trained with synthetic images generated by the TA-CycleGAN to segment the F-actin nanostructures on real live-cell STED images. The segmentation predictions generated by the U-Net*_Live_* (bottom-right) are similar to the manual expert annotations (top-right). **e,** The semantic segmentation task is used to train the TA-GAN*_Live_*. The generator of the TA-GAN*_Live_* takes as input the confocal image as well as a STED sub-region and a decision matrix indicating the position of the STED sub-region in the field of view (Methods). **f,** Real and synthetic live-cell STED images of F-actin generated with TA-GAN*_Live_*. The annotations of both real and synthetic images are obtained with the U-Net*_Live_*.

The TA-CycleGAN includes two generators that are trained to first perform a complete cycle between the two domains (fixed- and live-cell STED imaging), and then to compare the ground truth input image with the generated end-of-cycle image (Figure 3c). In the generic CycleGAN model, the losses are minimized when the generated images appear to belong to the target domain and the MSE between the input and output is minimized. In TA-CycleGAN we add a task network, here the U-Net*_F ixed−dend._*, which performs the semantic segmentation of dendritic rings and fibers. The U-Net*_F ixed−dend._* is applied to the real fixed STED images and the end-of-cycle reconstructed fixed STED images (Figure 3c). The generation loss is computed as the mean squared error between these segmentation masks. At inference, the trained TA-CycleGAN translates images of a given structure (here F-actin) but with image features (e.g. spatial resolution, signal-to-noise ratio, background level) corresponding to the target domain (here live-cell imaging). The *Translated F-actin dataset* was generated by applying the TA-CycleGAN to the *Dendritic F-actin dataset* (Supplementary Figure 11).

The *Translated F-actin dataset*, along with the expert annotations from the initial *Dendritic F-actin dataset*, is used to train the U-Net*_Live_* segmentation network to segment F-actin structures in images from the live-cell domain without requiring to annotate the *Live F-actin dataset* (Supplementary Figure 11, 3d). To confirm that training on synthetic domain-adapted images generalizes to real live-cell STED images, the AUROC was computed between the U-Net*_Live_* segmentation masks and manual ground truth annotations generated on 28 images by an expert in a user study (0.76 for rings and 0.83 for fibers, Extended data fig. 2, Supplementary Figure 12, and Supplementary Figure 13). In comparison, when applied to live-cell STED, the U-Net*_F ixed_* trained only on real images of fixed neurons achieves an AUROC of only 0.60 and 0.59 for the segmentation of rings and fibers respectively. Thus, domain adaptation with TA-CycleGAN enables the use of synthetic images to train a modality-specific segmentation network (here U-Net*_Live_*) when no real annotated dataset is available for training. This facilitates the cumbersome step in the training of any supervised machine learning method: creating data specific annotations. We next train TA-GAN*_Live_* for resolution enhancement of live F-actin confocal images using the *live F-actin dataset* and the pretrained U-Net*_Live_* as the auxiliary task network (Figure 3e, Methods). The annotations generated by the U-Net*_Live_* are used to compute the TL. Thus our image translation approach allows to train a TA-GAN to generate synthetic images from live-cell confocal images of F-actin in neurons (TA-GAN*_live_*) as well as a segmentation network adapted to the live-cell imaging domain (U-Net*_Live_*), without the need to annotate the *Live F-actin dataset* (Supplementary Figure 14).

### Imaging Assistance: Automated modality selection with TA-GAN

Optimizing light exposure is of particular concern for live-cell imaging, where multiple acquisitions over an extended period of time might be required to observe a dynamic process. In super-resolution microscopy, repeated imaging with high intensity illumination can cause photobleaching which quickly diminishes the signal quality (Supplementary Figure 15 and Extended data fig. 3). We evaluate how the integration of the TA-GAN*_Live_* in the acquisition loop of a STED microscope can guide imaging sequences for time-lapse live-cell microscopy. We apply our approach to detect the activity-dependent remodeling of dendritic F-actin from periodical rings into fibers in living neurons, which was previously observed in fixed neurons but could not be monitored in living neurons due to technical limitations [14]. For a given image acquisition sequence, we first acquire a confocal image (Figure 4a, Step 1). We next use a MC Dropout approach [47] to generate 10 possible synthetic STED images with the TA-GAN*_Live_*. We apply a different random dropout mask for each image generated (Figure 4a, Step 2). This use of MC Dropout with GANs has been previously demonstrated on natural images [48, 49] and serves as an estimation of the TA-GAN*_Live_*’s variability over the generated nanostructures. We next measure the optical flow between the 10 synthetic images (Figure 4a, Step 3, and Methods). The sub-region with the highest mean optical flow is acquired with the STED modality (Step 4) and given as an input to the TA-GAN*_Live_* together with the corresponding confocal image of the full field of view (Figure 4a, Step 5). This step helps minimizing the effect of signal variations encountered in live-cell imaging. The TA-GAN*_Live_* generates, with different dropout masks, 10 new synthetic images of the ROI, which are segmented by the U-Net*_Live_* to detect the presence of F-actin fibers (Figure 4a, Step 6). The segmentation predictions of the U-Net*_Live_* for the synthetic images are used to decide whether or not a real STED image should be acquired at a given time point (Figure 4a, Step 7). The acquisition of a complete frame using the STED modality is triggered when either 1) the segmentation prediction on the synthetic STED image is different from the one obtained on the last acquired real STED image (Figure 4b-e), or 2) there is high variability in the segmentation predictions on the 10 synthetic STED images (Figure 4f-i, Methods).

**Fig. 4:**
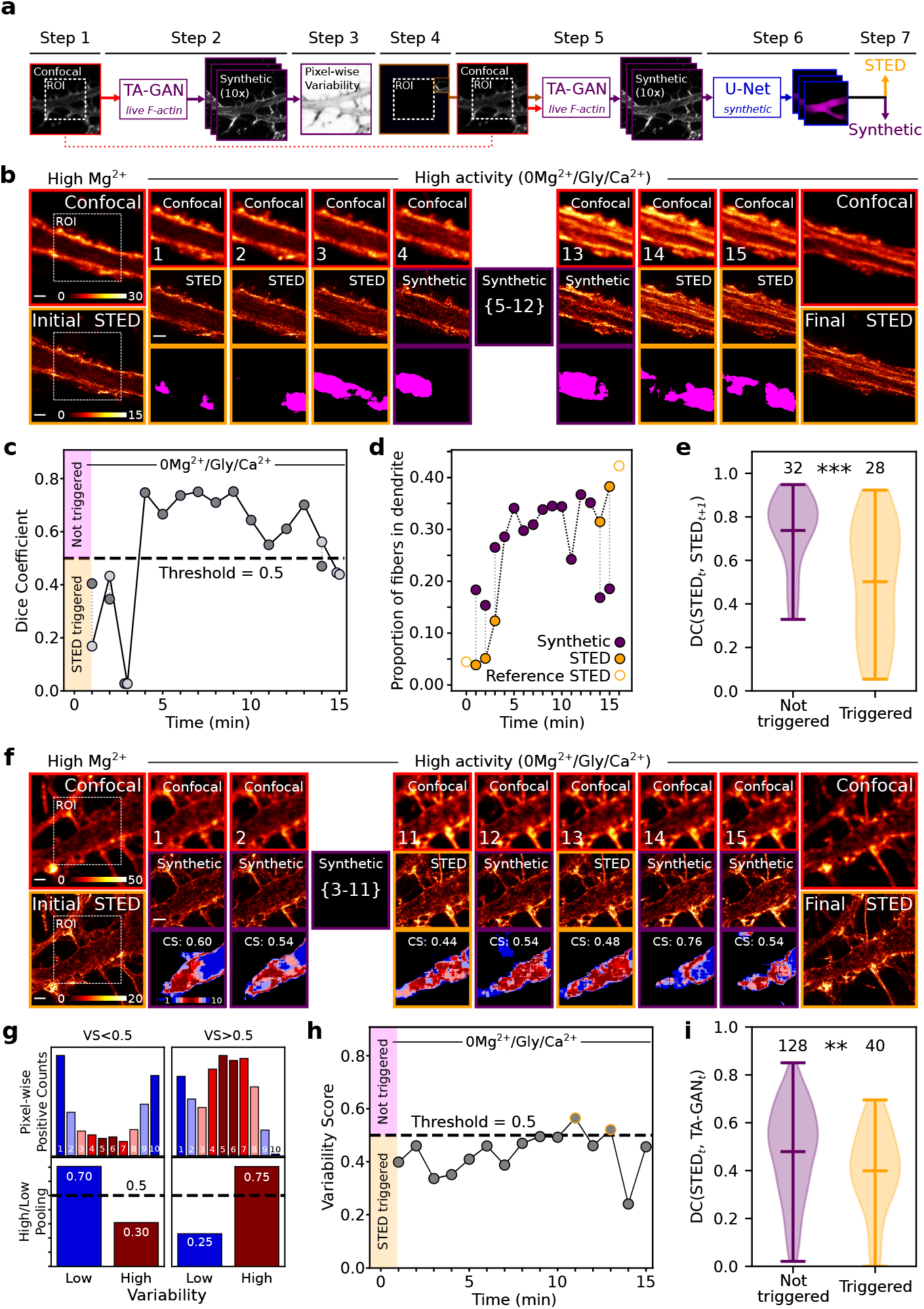
Live-cell imaging assistance with the TA-GAN*_Live_*. **a,** Step-by-step imaging assistance pipeline using the TA-GAN*_Live_* in the live-cell acquisition loop. **b,** Live-cell imaging of dendritic F-actin before (initial), during (frames 1-15) and after (final) application of a stimulation solution (0Mg^2+^/Gly/Ca^2+^). Shown are the confocal (red, top row), synthetic (purple, middle row), real STED images when acquired (orange, middle row), and corresponding segmentation masks for F-actin fibers (bottom row). **c,** The DC at each time point measured between the current synthetic image and the last acquired reference STED image for the sequence shown in **b**. Dark gray points indicate that the last acquired real STED (used as reference) is from a previous time step while light gray points indicate that a new STED is acquired at this time step, and the DC is recomputed with this new reference. **d,** Proportion of dendritic F-actin fibers at each time point segmented by the U-Net*_Live_* on either the real STED (orange) or the synthetic STED (purple) images. When a real STED acquisition is triggered, the proportion of fibers in both images is compared (dotted line). Initial and final reference STED images (empty orange bullets) are acquired at each round. **e,** The DC is computed for the F-actin fiber segmentation on control sequences of two consecutive real STED images (time points *t* and *t+1* )). The segmentation of the STED*t* image is used as reference and the DC is computed with the segmentation mask on the STED*_t_*_+1_ image. When a real STED image acquisition would not have been triggered by the threshold-based approach, the DC between the segmentation masks of the two real STED is higher. N=60 control sequences of two consecutive confocal-STED pairs. **f,** Live-cell imaging of dendritic F-actin using the same stimulation as in **b**. The TA-GAN*_Live_* variability maps are shown on the bottom row. **g,** Example histograms of the pixel-level positive counts over the segmentation of 10 synthetic images (top) and high/low variability pooling. On the left, the VS is below 0.5 (no trigger); on the right, the VS is above 0.5 (STED triggered). **h,** The VS at each time point for the sequence shown in **f** When the VS is above 0.5, the number of high variability pixels exceeds the number of low variability pixels (g, right), which triggers the acquisition of a real STED image (orange circles). **i,** The DC is computed between the segmentation masks of synthetic and real STED image from the same time point. When a STED acquisition would have been triggered using the VS criterion, the DC between the two corresponding images is lower. Statistical analysis: Mann-Whitney U test [50] for the null hypothesis that the two distributions are the same. (∗∗∗ *p <* 0.001, ∗∗ *p <* 0.01). Scale bars: 1 µm.

For the first acquisition scheme we calculate at each time point the mean DC between the segmentation masks from the 10 generated synthetic images and the last real STED image (Figure 4b, Supplementary Figure 16 and Supplementary Figure 17). A new STED image is acquired if the mean DC between the synthetic and the reference real STED images is below a predefined threshold of 0.5 (Figure 4c). Using paired confocal and real STED images acquired at the end of the imaging sequence (15 minutes), we show for the first time an increase in F-actin fibers proportion in living neurons (Figure 4b, last frame, Extended data fig. 4). Based on control sequences of two consecutive STED and confocal images pairs, we measure that the segmentation masks of those real STED images are more similar for sequences that would not have triggered a new real STED image acquisition, indicating that STED acquisitions are triggered at time points of higher biological change (Figure 4e and Supplementary Figure 18). The value of the DC threshold is chosen based on preliminary imaging trials and previous knowledge about the remodelling extent and dynamics, which, depending on the experimental context, is not always available prior to the imaging experiment.

We developed a second method to trigger STED image acquisitions, which is based on the variability in the predictions of the TA-GAN*_Live_*. This approach is particularly useful when not enough previous knowledge on the expected structural change is available to define a threshold for the DC prior to the experiment. With this acquisition scheme, for each confocal acquisition, we measure the pixel-wise variability of the segmentation predictions on the 10 generated synthetic STED images. Pixels predicted to belong to the same class (fibers or not fibers) in *≥*80 % of the synthetic images are defined as low variability pixels, and pixels predicted to belong to the same class in *<*80 % by the TA-GAN*_Live_* are defined as high variability pixels (Figure 4g). The proportion of high-variability pixels corresponds to the variability score (VS, Supplementary Figure 19). When the variability score is higher than 0.5 for the ROI, a full STED image is acquired (Figure 4f-i and Supplementary Figure 20). We validate the VS criterion on a set of real STED reference images and their corresponding synthetic counterparts. On these images we measure a higher DC between the segmentation masks, when a real STED image acquisition would not have been triggered by the VS threshold (Figure 4i and Supplementary Figure 18). This indicates that the VS is a good indicator of the similarity between the real and synthetic STED image at a given time point. This approach can be beneficial to detect unexpected patterns and rare events. For both approaches, modulation of the STED modality acquisition frequency can be achieved by adapting the DC or VS thresholds.

## Discussion

We introduce TA-GAN for resolution enhancement and domain adaptation. We demonstrate its applicability to optical nanoscopy (Extended data fig. 5) and show that an auxiliary task assisting the training of a generative network improves the reconstruction accuracy of nanoscopic structures. The applicability of our method is demonstrated for paired confocal and STED microscopy datasets of F-actin in axons and dendrites, synaptic protein clusters, simulated nanodomains as well as for paired brightfield and SIM images of dividing S. aureus bacterial cells. We show that the TA-GAN method is flexible and can be trained with different auxiliary tasks such as binary segmentation, semantic segmentation, and localization. For unpaired datasets, we introduce the TA-CycleGAN model and demonstrate how the structure preserving domain adaptation opens up the possibility to create paired datasets of annotated images that cannot be acquired simultaneously. Those synthetic STED images from the live-cell domain can be used to train a neural network that performs well for the segmentation of F-actin nanostructures in *real* STED images, without the need for manual re-annotations of the new live-cell imaging dataset. The TA-GAN for resolution enhancement in living neurons can be integrated into the acquisition loop of a STED microscope (Figure 4). We validate how this TA-GAN model can be helpful in assisting a microscopists by automating decisions in live-cell optical nanoscopy acquisition sequences. The TA-GAN increases the informative value of each confocal acquisition and automatically triggers the acquisition of a STED image only in the regions and time-steps where this acquisition is informative enough due to variations or uncertainties in the predicted nanostructures (Figure 4).

Future work in calibrating the network’s probabilistic output could lead to an improved quantification of its confidence. Multiple successive frames could also be given as input to the generator to introduce temporal information instead of using static frames individually. This could enable the generator to decode the rate of biological change and introduce this knowledge to the next frame prediction, leading to smoother transitions between synthetic images. The TA-GAN model, as presented here, enables the visualization of biological dynamics over longer sequences with reduced photobleaching effects. Thus, TA-GAN-assisted STED nanoscopy can guide microscopists for optimized acquisition schemes and reduced light exposure.

## Acknowledgments

Francine Nault and Sarah Pensivy for neuronal cell culture. Gabriel Leclerc for the FIJI macro for segmentation. Annette Schwerdtfeger for proofreading the manuscript. Funding was provided by grants from the Natural Sciences and Engineering Research Council of Canada (NSERC) (F.L.C. and C.G.), the Canadian Institute for Health Research (CIHR) (F.L.C.), and the Neuronex Initiative (National Science Foundation 2014862, Fond de recherche du Québec - Santé) (F.L.C.). C.G. is a CIFAR Canada AI Chair and F.L.C. is a Canada Research Chair Tier II. C.B. is supported by scholarships from NSERC, from the Fonds de Recherche Nature et Technologie (FRQNT) Quebec, from the FRQNT strategic cluster UNIQUE, and by a Leadership and Scientific Engagement Award from Université Laval. T.W. was supported by a postdoctoral scholarship from the FRQNT strategic cluster UNIQUE. A.B. is supported by scholarships from NSERC, FRQNT and from the FRQNT strategic cluster UNIQUE.

## Declarations

### Availability of data and materials

The *S. aureus dataset* from Spahn et al. [44] is available at https://zenodo.org/ record/5550933%23.Y6IhFNLMJH4 and https://zenodo.org/record/5551141#.Y6IjBdLMJH5. All other datasets are available at https://s3.valeria.science/ flclab-tagan/index.html.

### Code availability

The codes, trained models, and sample images needed to test the TA-GAN architecture are available at https://github.com/FLClab/TA-GAN, as well as instructions on how to adapt the dataloaders and train the networks for new experiments.

### Author contributions

C.B., F.L.C. and C.G. designed the method. C.B. performed all deep learning experiments, implemented the live imaging automatic acquisitions, and analysed the results. A.B. and B.T. generated the simulated nanodomain dataset. A.D. and T.W. performed the live imaging experiments. A.D. contributed to the integration of the TA-GAN on the microscope. A.B. provided trained networks, helped manage the data, analysed results on the simulated nanodomain dataset, and designed the website. C.B., F.L.C., and C.G. wrote the manuscript.

## Methods

### Sample preparation and STED microscopy

#### Cell culture

Dissociated rat hippocampal neurons were prepared as described previously [14, 51] in accordance with and approved by the animal care committee of Université Laval. For live-cell STED imaging, the dissociated cells were plated on PDL-Laminin coated glass coverslips (18 mm) at a density of 322 cells/mm^2^ and used at DIV 12-16.

#### STED microscopy

Live-cell super-resolution imaging was performed on a 4-color Abberior STED microscope (Abberior Instruments, Germany) using a 40 MHz pulsed 640 nm excitation laser, a ET685/70 (Chroma, USA) fluorescence filter, and a 775 nm pulsed (40 MHz) depletion laser. Scanning was conducted using a pixel dwell time of 5 µs, a pixel size of 20 nm, and 8 line repetition sequence. The STED microscope was equipped with a motorized stage and auto-focus unit. The imaging parameters used are described in Table 1.

**Table 1:** Imaging parameters used for the live-cell experiments and compared with the parameters used in [14] for F-actin imaging in fixed neurons. The laser power was measured at the back focal plane.

|  | Imaging parameters<br>for the generation of the<br>*Dendritic F-actin dataset* from [14] | Imaging parameters<br>for the generation of the<br>*Live F-actin dataset* |
| --- | --- | --- |
| **Fluorophore** | Phalloidin-STAR635 | SiR-Actin |
| **Excitation power)** | 6 μW | 6.3 μW |
| **Depletion power (at BFP)** | 180 mW | 110 mW |
| **Pixel Dwelltime** | 15 μs | 5 μs |
| **Line steps** | 4 | 8 |
| **Pixel size** | 20 nm | 20 nm |

The cultured neurons were pre-incubated in HEPES buffered artificial cerebrospinal fluid (aCSF) at 33*^◦^*C with SiR-Actin (0.5 µM, SpiroChrome) for 8 minutes and washed once gently in SiR-Actin-free media. Imaging was performed in HEPES buffered aCSF of high Mg^2+^/low Ca^2+^ (in mM: NaCl 98, KCl 5, HEPES 10, CaCl_2_ 0.6, Glucose 10, MgCl_2_ 5) using a gravity driven perfusion system. Neuronal stimulation was performed with an HEPES buffered aCSF containing 2.4 mM Ca^2+^, Glycine and without Mg^2+^ (in mM: NaCl 98, KCl 5, HEPES 10, Glycine 0.2, CaCl_2_ 2.4, Glucose 10). Solutions were adjusted to an osmolality of 240 mOsm per kg and a pH of 7.3.

### Datasets

#### Axonal F-actin dataset

The publicly available *Axonal F-actin dataset* [14] was used to train the TA-GAN*_Ax._* for confocal-to-STED resolution enhancement of axonal F-actin nanostructures using the binary segmentation of F-actin rings as the auxilliary task. The original dataset consisted of 516 paired confocal and STED images (224 *×* 224 pixels, 20 nm pixel size) of axonal F-actin in fixed cultured hippocampal neurons from Lavoie-Cardinal *et al.* [14]. 31 images from the original dataset were discarded for not containing annotated axonal F-actin rings. The remaining images were split into a training set (377 images), a validation set (56 images), and a testing set (52 images). The manual polygonal bounding box annotations of the axonal F-actin periodical lattice (F-actin rings) from the original dataset were retained (Figure 1b).

#### Dendritic F-actin dataset

The publicly available *Dendritic F-actin dataset* was used to train the TAGAN*_Dend._* for confocal-to-STED resolution enhancement of dendritic F-actin nanostructures using the semantic segmentation of F-actin rings and fibers as the auxiliary task. The *Dendritic F-actin dataset* was also used to train the TA-CycleGAN for domain adaptation. The original dataset from Lavoie-Cardinal *et al.* [14] was split into a training set (304 images), a validation set (54 images), and a testing set (26 images, 12 for low activity and 14 for high activity). The dataset consists in paired confocal and STED images of the dendritic F-actin cytoskeleton in fixed cultured hippocampal neurons, which had been manually annotated using polygonal bounding boxes. The training and validation crops were taken from large STED images (between 500 *×* 500 and 3000 *×* 3000 pixels, 20 nm pixel size) using a sliding window of size 224 *×* 224 pixels with no overlap. If less than 1 % of the pixels of the crop were annotated as containing a structure of interest (F-actin rings and/or fibers), the crop was discarded from the set. This operation resulted in 4,331 crops for training and 659 crops for validation.

#### Simulated nanodomains dataset

We used the pySTED image simulation platform [39] to create a simulated dataset of nanodomains within a dendritic spine. The pySTED simulator requires as input a matrix providing the position and number of fluorescent molecules for each pixel in the field of view, referred to as a datamap. Each datamap (64 *×* 64 pixels or 1.28 *×* 1.28 µm) consisted of a mushroom spine-like shape (between 0.12 and 0.48 µm^2^) containing N (1-6) regions (20 *×* 20 nm) with a higher fluorophore concentration, which we refer to as nanodomains. In the majority of the images, the simulated STED modality was required to resolve all nanodomains. The position of the nanodomains was randomly distributed on the edge of the synapse (*<* 140 nm away from the edge) with a minimal distance of 40 nm between nanodomains. We allowed random rotation and translation of the spine making sure that the nanodomains were kept within the field of view. For training, we generated a total of 1200 simulated datamaps (200 for each number of nanodomains). The training and validation datasets were split using a 90-10 ratio. The localization maps are matrices of size 64 *×* 64 pixels, where the value of each pixel is the cubic root of the distance to the closest nanodomain. Two testing datasets were created. The first one consisted in 75 simulated datamaps with different numbers of nanodomains (2-6). The second one consisted in 80 images with two nanodomains, where the distance between the pair of nanodomains varies from 40 nm to 450 nm.

#### Synaptic protein dataset

The publicly available *Synaptic protein dataset* consists of paired two-color STED and confocal images of the synaptic protein pair PSD95 (postsynapse) and Bassoon (presynapse) in fixed hippocampal neurons obtained from Wiesner *et. al* [41]. The dataset was split into a training set (32 images), a validation set (2 images) and a testing set (10 images). The confocal and STED images from the training and validation sets were first registered using the pipeline presented in Supplementary Figure 21, resulting in 690 crops for training and 35 crops for validation. The segmentation maps were generated by automatically segmenting the STED images using wavelet transform decomposition [42] with the same parameters (scales 3 and 4) as in Wiesner *et al.* [41]. No segmented clusters were discarded based on size or position, following the intuition that even the smallest structures should be generated. The localization maps were created from a black image by placing a white pixel at the position of the intensity-weighted centroid of each segmented cluster, and then applying a Gaussian filter with a standard deviation of 2 (Supplementary Figure 22).

#### S. aureus dataset

We used the brightfield images and the corresponding structured illumination microscopy (SIM) images from the publicly available *S. aureus* dataset for segmentation from Spahn *et al.* [44]. This dataset includes 12 images (6 for training, 1 for validation, 5 for testing) with manual whole-cell annotations. The brightfield images (80 nm/px) were rescaled to the size of the SIM images (40 nm/px) using bilinear interpolation, and the cell annotations were rescaled using nearest-neighbor interpolation. The whole-cell annotations were converted to binary segmentation maps with pixel values of 0 for background and 1 for cells. Those whole-cell segmentation maps were used as low-resolution annotations. High-resolution annotations highlighting the cell division boundary were generated from the SIM images. To generate the high-resolution annotations we first applied a Sobel filter to the SIM images to find the outer and inner edges of the cells. We next applied a threshold corresponding to 20 % of the maximum value of the filtered result. This resulted in a mask of the boundary between dividing cells as well as of the cell outer membrane. We applied a Sobel filter to the low-resolution annotations with a threshold of 0 followed by a Gaussian filter with a standard deviation of 1 to generate a binarized cell border mask. The binarized cell border mask was subtracted from the mask of the outer and inner cell borders to generate the final high-resolution annotations.

The training crops were generated using a sliding window of size 256 *×* 256 pixels with an overlap of 128 pixels. Crops were discarded if they contained less than 3 % annotated pixels. The validation crops were generated using the same sliding window method, but for a size of 128 x 128 pixels without overlap. All validation crops were considered, regardless of the percentage of annotated pixels. The resulting dataset comprised 202 training crops and 64 validation crops.

#### Live F-actin dataset

The *Live F-actin dataset* was acquired for this study and was used to train: 1) the TA-CycleGAN for live and fixed domain adaptation, and 2) the TAGAN*_Live_*. The *Live F-actin dataset* consists in 904 paired STED and confocal images of F-actin stained with the fluorogenic dye SiR-Actin (Spirochrome, US) in living hippocampal cultured neurons (see Table 1). The dataset was split into a training set (833 images) and a validation set (71 images). The images were of variable size (from a minimum width of 2.76 to a maximum of 49.1 µm, pixel size is always 20 nm).

#### Translated F-actin dataset

The *Translated F-actin dataset* was used to train the TA-GAN*_Live_*. This dataset corresponds to the *Dendritic F-actin dataset* adapted to the live-cell STED imaging domain using the TA-CycleGAN for fixed-to-live domain adaptation. It contains the same number of images, the same train/valid/test splits, and the same image characteristics (crop size, pixel size, annotations) as the *Dendritic F-actin dataset*.

### TA-GAN training procedure

The TA-GAN was developed from the conditional GAN (cGAN) model for image-to-image translation pix2pix [35], available at https://github.com/ junyanz/pytorch-CycleGAN-and-pix2pix. Comparable methods using cGANs for enhancing the resolution of microscopy images are trained using pixel-wise generation losses to compare the generatd image with the ground truth, such as mean squared error (MSE) [5], absolute error [15, 16] or structural similarity index [17, 18]. For the TA-GAN, the generation loss is computed by comparing the output of an auxiliary task network applied on the real (ground truth) and generated (synthetic) images (Figure 1a). The other standard losses for conditional GANs[35] are also used for TA-GAN : the discrimination losses for the classification of the real and generated images, and the GAN loss for the misclassification of generated images. The networks (generator, discriminator and task network) are optimized using the Adam optimizer with momentum parameters *β*_1_ = 0.5, *β*_2_ = 0.999 for all TA-GAN models. We follow the same approach as the pix2pix paper[35]: at each epoch we alternate between one gradient descent step on the discriminator, then one step on the generator, then one step on the task network. Table 2 summarizes the settings for the resolution enhancement experiments presented in this paper, and Table 3 presents the hyperparameters used for training the TA-GAN for each of these experiments.

**Table 2:** Summary of experimental settings using the TA-GAN for microscopy resolution enhancement.

|  | Dataset | Input | Output | Task | Annotations | Task network |
| --- | --- | --- | --- | --- | --- | --- |
| TA-GAN<br>*Ax.* | Axonal F-actin | Confocal (1 channel) | STED (1 channel) | Segmentation of axonal F-actin rings | Bounding boxes (1 class) | Not pretrained |
| TA-GAN<br>*Nano.* | Simulated nanodomains | Simulated confocal (1 channel) | Simulated STED (1 channel) | Localization of nanodomains | Ground truth positions | Not pretrained |
| TA-GAN<br>*Syn.* | Synaptic Proteins (localization) | Confocal (2 channels) | STED (2 channels) | Localization of synaptic protein cluster centroids | Wavelet detected centroids (2 channels) | Not pretrained |
| TA-GAN<br>*Syn.* | Synaptic Proteins (segmentation) | Confocal (2 channels) | STED (2 channels) | Segmentation of protein clusters | Wavelet segmented clusters (2 channels) | Not pretrained |
| TA-GAN<br>*SA.* | S. aureus (LR labels) | Bright-field (1 channel) | SIM (1 channel) | Segmentation of individual cells | Circular segmentation masks (1 class) | Not pretrained |
| TA-GAN<br>*SA.* | S. aureus (HR labels) | Bright-field (1 channel) | SIM (1 channel) | Segmentation of cell division lines | Segmentation masks created from SIM images (1 class) | Not pretrained |
| TA-GAN<br>*Dend.* | Dendritic F-actin | Confocal (1 channel) | STED (1 channel) | Semantic segmentation of dendritic F-actin rings and fibers | Bounding boxes (2 classes) | Not pretrained |
| TA-GAN<br>*Live* | Live F-actin | Confocal + decision matrix + STED sub-region | STED (1 channel) | Semantic segmentation of dendritic F-actin rings and fibers | None | Pretrained and frozen |

**Table 3:** Hyperparameters used to train the TA-GAN model on the four datasets presented.

| Hyperparam. | TA-GAN<br>*Ax.* | TA-GAN<br>*Nano.* | TA-GAN<br>*Syn.* | TA-GAN<br>*SA.* | TA-GAN<br>*Dend.* | TA-GAN<br>*Live* |
| --- | --- | --- | --- | --- | --- | --- |
| # train. images | 377 | 1,080 | 1,841 | 202 | 4,331 | 753 |
| # valid. images | 56 | 120 | 509 | 64 | 659 | 47 |
| Image size | 224x224 | 64x64 | 512x512 | 256x256 | 224x224 | Variable |
| Generator | 9-blocks ResNet [52] |  |  |  |  |  |
| Task net. | 6-blocks ResNet |  |  |  |  | U-Net128 [53] |
| Discriminator | PatchGAN [54] |  |  |  |  |  |
| Batch size | 8 | 256 | 32 |  | 32 | 16 |
| Learn. rate | 0.0002 |  |  |  |  |  |
| Weight GAN | 1 |  |  |  |  |  |
| Weight GEN | 10 | 10 | 10 |  | 1 | 10 |
| Data augm. | Horizontal and vertical flips, random crops, 90 degrees rotations |  |  |  |  |  |
| Crop size | 128 | 64 | 128 |  | 128 | 256 |
| # epochs | 1000 | 200 |  |  | 500 | 5000 |

### TA-GAN training with segmentation auxiliary tasks

The TA-GAN*_Ax._*, TA-GAN*_Dend._*, TA-GAN*_Syn._*, and TA-GAN*_SA._*were trained for resolution enhancement using the segmentation of sub-diffraction biological structures as the auxiliary task. The output of the segmentation network was compared with the ground truth annotations using a MSE loss. The loss computed from the real STED image (Task Loss, TL in Figure 1a) was back- propagated to the segmentation network to optimize its weights, and the loss from the synthetic STED image (GEN) optimizes the generator. The other losses computed were standard cGAN losses: the GAN loss (GAN: misclassification of synthetic images as real images), the discriminator losses (DR: classification of real images as real, and DG: classification of generated images as synthetic). The validation losses were not used for early-stopping because of the adversarial nature of GANs. The validation images were instead used as a qualitative assessment of the training progress to select the best iteration for testing the model.

For the TA-GAN*_Ax._*, the auxiliary task was the segmentation of axonal F-actin rings. The output of the auxiliary task network was the predicted segmentation maps of F-actin rings. The spatial resolution of the real and synthetic images were not significantly different (Supplementary Table 1, Supplementary Figure 23).

For the TA-GAN*_Dend._*, the auxiliary task was the semantic segmentation of dendritic F-actin rings and fibers. The output of the auxiliary task network was a two-channel image, with the predicted segmentation maps of F-actin rings in the first channel and of F-actin fibers in the second channel. The spatial resolution of the real and synthetic images were not significantly different (Supplementary Table 1, Supplementary Figure 23).

For the TA-GAN*_SA._*, the auxiliary task was either whole-cell segmentation (LR annotations) or the segmentation of the boundary between dividing cells (SR annotations). The output of the auxiliary task network is a one-channel image with the predicted segmentation maps of either whole cells or the dividing cell boundaries, respectively.

For the TA-GAN*_Syn._* trained using a segmentation task, the output of the segmentation network is a two-channel image with the predicted segmentation maps of PSD95 clusters in the first channel and Bassoon in the second channel. The spatial resolution of the real and synthetic images were not significantly different (Supplementary Table 1, Supplementary Figure 23).

### TA-GAN training with a localization auxiliary task

The TA-GAN*_Syn._* and TA-GAN*_Nano._*for confocal-to-STED resolution enhancement were trained using a localization network to compute the generation loss. The localization network took a STED image as input to output a map of dots indicating the intensity-weighted centroids of all detected clusters in the STED image.

The TA-GAN*_Syn._* was trained on the *Synaptic protein dataset* using the two-channel confocal image rescaled and registered to the STED image. The generation loss (GEN in Figure 1) was the MSE between the weighted centroids of the real STED image and the localization predictions from the task network on the synthetic image. The spatial resolution of the real and synthetic images were not significantly different (Supplementary Table 1, Supplementary Figure 23).

TA-GAN*_Nano._* was trained on the *Simulated nanodomain dataset* using the simulated confocal image as input. The generation loss was the MSE between the localization maps from the ground truth datamaps and the localization predictions from the task network on the synthetic image.

### TA-CycleGAN training for domain adaptation

The TA-CycleGAN model was developed from the CycleGAN model [35]. As for the standard CycleGAN, the TA-CycleGAN consists of 4 networks : two generators (one that translates the domain of fixed-cell STED imaging (*F* ) into domain of live-cell STED imaging (*L*), the other which translates domain *F* into domain *L*), and two discriminators (one for domain *F* , the other for domain *L*), which are combined with a fifth network : the auxiliary task network (Figure 3c). The TA-CycleGAN was applied to non-paired images, where the prediction of the generator for a given input cannot be compared to a corresponding ground truth. Instead, the generated synthetic image was passed through a second generator and converted back to the input domain where it was compared to the initial image (ground truth) for the computation of losses. The TA-CycleGAN for fixed-to-live domain adaptation was trained using two datasets: the *Dendritic F-actin dataset* (*F* ) and the *Live F-actin dataset* (*L*). The auxiliary task was the semantic segmentation or F-actin rings and fibers on the *Dendritic F-actin dataset*, for which manual bounding box annotations were available [14]. The U-Net*_F ixed−dend._* was already optimized for the semantic segmentation of F-actin rings and fibers in fixed-cell STED images [14]. The generation loss was the MSE between the U-Net*_F ixed−dend._* segmentation prediction on the real fixed cell image (*Fixed* ), and the end-of-cycle fixed cell image (*Fixed_rec_*) (3c).

### Training procedures of resolution enhancement and denoising baselines

#### Enhanced Super-Resolution Generative Adversarial Networks (ESRGAN x4)

ESRGAN x4 (Enhanced Super-Resolution Generative Adversarial Networks) [33] is a state-of-the-art method for upsampling natural images. ESRGAN was implemented from the public GitHub repository (https://github.com/xinntao/Real-ESRGAN). We fine-tuned ESRGAN on two of our datasets, the *Axonal F-actin dataset* and the *Simulated nanodomains dataset*, using the code and pretrained weights released with the most recent iteration of the model, Real-ESRGAN[34]. For both datasets, the input of the model is the confocal image, and the target output is the corresponding STED image upsampled 4 times using nearest neighbor interpolation. Even though the confocal and STED images are the same size, the upsampling had to be kept in the model in order to use the pretrained weights. ESRGAN was fine-tuned for 50,000 iterations. The model was applied to the validation images to ensure training had converged after 50,000 iterations. All default parameters proposed by the authors were used, except for the input crop size and batch size (128 pixels and 4 for the *Axonal F-actin dataset*, 64 pixels and 16 for the *Simulated nanodomains dataset* ).

#### Content-Aware image REstoration (CARE)

Content-Aware image REstoration (CARE) [16] uses a U-Net for deblurring, denoising, and enhancing fluorescence microscopy images. CARE was implemented from the public GitHub repository (https://github.com/CSBDeep/ CSBDeep). We used the standard CARE network for image restoration and enhancement. The residual U-Net generator was optimized from scratch on our datasets. The original CARE model does not use data augmentation, since it’s trained on unlimited simulated images. We augmented our datasets prior to training the CARE models so that the number of training images is similar to the one used for the original model trained on simulated images (8,000 synthetic pairs of 128 x 128 pixels). For the *Axonal F-actin dataset*, each image from the training set is augmented 32 times by cropping the four corners into 128x128 crops and applying the 8 possible flips and rotations to each corner crops. The 377 224 x 224 images were augmented into 12,064 different crops. For the *Simulated nanodomains dataset*, the 64 x 64 images were too small to be further cropped, but were instead augmented 8 times using flips and rotations. The 1,080 training images were augmented into 8,640 different images. The patience parameter for the learning rate decay function was adjusted from 10 to 20 epochs after noticing that the learning rate was reduced too abruptly to allow the training loss to properly converge. Except for the patience of the learning rate decay function, default hyperparameters were used and the model was trained for 100 epochs using a mean absolute error loss. The epoch that reached the lowest validation loss was used for testing.

#### Residual channel attention networks (RCAN)

RCAN [55] uses residual channel attention networks to increase the resolution of natural images. 3D-RCAN [15] adapts the original model to denoise and sharpen fluorescence microscopy image volumes. We used the code implemented with Tensorflow and Keras from the publicly available GitHub repository (https://github.com/AiviaCommunity/3D-RCAN). We used the same patch size as for training the TA-GAN*_Ax._* (128 x 128 pixels) and the TA-GAN*_Nano._* (64 x 64 pixels). We trained different RCAN models using configurations of hyperparameters that were inspired by both the 2D [55] and the 3D [15] versions. We first trained a model on the *Axonal F-actin dataset* with the hyperparameters from the 2D RCAN version. Even though both training and validation losses had converged, the output obtained with the weights from the epoch of lowest validation loss (epoch 205 out of 1000) is an unrecognizable and smoothed version of the input. We hypothesize that this version of the model is too deep (15M trainable parameters) for the number of training images. We trained a second version of RCAN using the hyperparameters from Chen *et al.* [15]. The loss when training this model quickly converges to a minimum (epoch 34 out of 300), and the resulting images are smoothed versions of the input confocal image. This simplified version of RCAN might be too lightened for the 2D context. The architecture that ended up performing the best with our datasets mixes hyper-parameters from both implementations.

1) We used 2D convolutions because our images are 2D, as in RCAN. 2) We set the number of residual groups to 10 in the residual in residual structure, as in 3D-RCAN. 3) The residual channel attention blocks was set to 20, as in RCAN. 4) We set the number of convolution layers in the shallow feature extraction and residual in residual structure to 32, as in 3D-RCAN. 5) We set the reduction ratio to 8 as in 3D-RCAN. 6) The upscaling module was removed because the confocal and STED images are the same size, as is the case for 3D-RCAN. This RCAN model was trained for 1000 epochs for both datasets to ensure convergence of the validation loss. The model reaching the lowest validation loss (epoch 838 for the *Simulated nanodomains dataset*, epoch 398 for the *Axonal F-actin dataset* ) was used for testing.

#### Conditional GAN for image-to-image translation (pix2pix)

Pix2pix [35] is a state-of-the-art method for image-to-image translation in natural images. It was implemented with Pytorch from the publicly available GitHub repository (https://github.com/junyanz/ pytorch-CycleGAN-and-pix2pix). The TA-GAN and pix2pix share the same architecture with or without the task assistance. For each experiment, the same hyperparameters and datasets as for the TA-GAN were used for training (Table 3), replacing only the generation loss with a pixel-wise MSE loss between the ground truth and generated STED images. The results from this baseline are compared to the TA-GAN for all fixed-cell datasets.

#### Denoising convolutional neural networks (DnCNN)

DnCNN (denoising convolutional neural networks) [36] is a state-of-the-art denoising method for natural images. The trained version of DnCNN [36] available at https://github.com/yinhaoz/denoising-fluorescence was directly applied to our test images for all datasets (Supplementary Figure 1). The datasets used in this study do not provide the required characteristics to retrain DnCNN (i.e., lack of images with different noise levels), therefore a published version of the DnCNN trained on the fluorescence microscopy denoising dataset [37] was used as is. It was included as a baseline to show how the confocal-to-STED and BF-to-SIM transformations are not denoising tasks.

#### Noise2Noise (N2N)

Noise2Noise[38] is a state-of-the-art deep learning denoising method that does not require clean (denoised) data for training. Like DnCNN, we used the training version available at https://github.com/yinhaoz/denoising-fluorescence and directly applied it to the test images from our datasets, without retraining or fine-tuning (Supplementary Figure 1).

### Evaluation of networks performance

#### Segmentation of F-actin nanostructures in synthetic STED images

The performance of the TA-GAN*_Ax._* was measured on the test set of the *Axonal F-actin dataset*. The MSE, PSNR, and SSIM were computed between the ground truth and synthetic STED images of the test set (Figure Extended data fig. 1). Additionally, U-Net*_F ixed−ax._*, a U-Net that was trained to segment axonal F-actin rings on real STED images only [14] (available at https://github.com/FLClab/STEDActinFCN) was used to produce segmentation masks of axonal F-Actin rings on the real and synthetic STED image pairs (Figure Extended data fig. 1 and Figure 3), which were compared using the Dice Coefficient (DC) and Intersection over Union (IOU) metrics. We used the trained weights provided and did not retrain U-Net*_F ixed−ax._* specifically for this work.

The performance of the TA-GAN*_Dend._* was evaluated on the test set of the *Dendritic F-actin dataset*. The U-Net*_F ixed−dend._*, a U-Net that was trained for the semantic segmentation of dendritic F-actin rings and fibers on real STED images only [14] (available at https://github.com/FLClab/STEDActinFCN), was used to segment the real and synthetic STED images. The segmentation masks of both F-actin rings and fibers were compared on the real and synthetic STED image pairs (Supplementary Figure 10). We used the trained weights provided and did not retrain U-Net*_F ixed−dend._* specifically for this work.

#### Assessment of synaptic protein cluster morphology

The perimeter, eccentricity, area, distance to nearest neighbor from the same channel, and distance to nearest neighbor from the other channel of the protein clusters from the *Synaptic protein dataset* were measured in the confocal, STED images and synthetic images (Figure 2, Supplementary Figure 7). The distribution of each morphological feature over all associated clusters from the test set images was computed using a Python library for Statistical Object Distance Analysis (pySODA) [41] (Supplementary Figure 7). A foreground mask was generated following Wiesner *et al.* [41]: applying a Gaussian blur (standard deviation of 10) on the sum of both STED channels, and thresholding the image using 50% of the mean intensity value. Only clusters from the foreground mask were considered for the analysis. The same parameters as in Wiesner *et al.* [41], which were optimized for real STED images of synaptic protein clusters, were used for the analysis: wavelet segmentation scales of 3 and 4, a minimum cluster area of 5 pixels, and minimum cluster width/height of 3 pixels. The weighted centroids of the detected clusters were calculated on the raw STED images.

#### Classification of S. Aureus cells

The TA-GAN*_SA._* performance was evaluated using the classification of dividing bacterial cells, which is a task that cannot be achieved using only the brightfield images. A simple threshold optimization applied on bright-field images was not sufficient to classify the cells as dividing or not (Supplementary Figure 9). A dividing bacterial cell is defined as having a clear boundary between the two dividing cells that can be identified in the SIM image. We trained the ResNet*_SA._* using the SIM images (training set) and the high-resolution annotations, to segment the dividing cell boundaries. The ResNet_SA._ is a ResNet-9 architecture trained for 200 epochs using a MSE loss, a learning rate of 0.0002, and Adam optimizer. All real SIM images and synthetic SIM images generated from pix2pix_SA._, TA-GAN_SA._ trained with low-resolution annotations, and TA-GAN_SA._ trained with high-resolution annotations are segmented by ResNet_SA._.

The dividing / non-dividing cells classification was based on the segmentation of the ResNet*_SA._*: 1) dividing if the segmentation mask contained at least 20 positive pixels, 2) non-dividing if the segmentation mask was empty for a given cell. For segmentation masks containing 1-19 pixels, the cells were identified as ambiguous and discarded. On the real SIM images test set, 251 cells were identified as non-dividing (single cells) and 159 as dividing (showing a clear boundary between the dividing cells).

#### User-study for the segmentation of live F-actin images

A set of 28 STED images (224 x 224 pixels) from the *Live F-actin dataset* test set was labeled by an expert using a FIJI [56] macro to test the performance of the U-Net*_Live_* trained on the *Domain adapted dendritic F-actin dataset* for the segementation of real live-cell STED images. In addition, a second set of 28 synthetic images, selected from the *Domain adapted dendritic F-actin dataset* was included in the user-study. The expert was presented with an image from one of the two sets, without being informed whether the image was real or synthetic. For each image, the expert draws polygonal bounding boxes that enclosed all regions identified as F-actin rings and fibers.

#### User-study for the localization of nanodomains

The positions of the nanodomains in the real and synthetic test images of the *Simulated nanodomains dataset* were identified by an expert to compare the localization performance of the TA-GAN*_Nano._* with the baseline methods.

The expert was presented with an image without being informed whether the image was real or synthetic, or by which baseline it was generated. For each image, the expert selects the pixel identified as the center of each nanodomain detected. To compute the F1-score, a detection is defined as a true positive if it is within 3 pixels of the ground truth position of a nanodomain center.

### TA-GAN-assisted live-cell STED microscopy

#### Training of the TA-GAN*_LiveF Actin_*

The TA-GAN*_Live_* for resolution enhancement of live-cell STED imaging was trained on the new and not previously annotated *Live F-actin dataset*. The auxiliary task was the semantic segmentation of dendritic F-actin rings and fibers. The original *Live F-actin dataset* did not include any manual annotations. To circumvent this limitation, the U-Net*_Live_* segmentation network was pretrained on the *Domain adapted dendritic F-actin dataset*. The pretrained U-Net*_Live_* was frozen during the TA-GAN*_Live_* training and was used to compute the MSE generation loss between the segmentation prediction of the real and the synthetic STED images.

To better adapt to cell-to-cell signal variations and experimental variability in live-cell STED images, the input of the generator has three channels: 1) the confocal image, 2) a real STED sub-region acquired in the vicinity of the region of interest (ROI), and 3) an image indicating the position of the STED sub-region (Figure 3e). Training using this three-channel input enables the generator to learn features from the STED sub-region and turns the resolution enhancement task into an image completion task.

#### Training of the U-Net*_Live_*

The U-Net*_Live_* was built around a U-Net-128 [53] architecture with batch normalization and two output channels (F-actin rings and fibers) for the segmentation of F-actin nanostructures in living-neurons.

The training of the U-Net*_Live_* required an annotated dataset of images of the live-cell domain. A random subset (2,069 training crops and 277 validation crops) of the *Dendritic F-actin dataset* was translated into the live-cell domain using the generator*_live_*(Figure Supplementary Figure 11). This resulted into the *Domain-adapted F-actin dataset*. The manual annotation from the fixed cell images were associated with the corresponding synthetic images from the live-cell domain (Figure Supplementary Figure 11a).

Random crops of 128 x 128 pixels of the *Domain-adapted F-actin dataset* and their corresponding annotations were used to train U-Net*_Live_* on images of the live-cell domain. Horizontal and vertical flips were used for data augmentation. Due to class imbalance in the training set, the segmentation loss for fibers was weighted by a factor of 2.5, which reflected the ratio of total annotated pixels for each class. The U-Net*_Live_* was trained for 1000 epochs and the iteration with the lowest segmentation loss over the validation set was kept for further use and testing. The optimal threshold to binarize the segmentation prediction was determined as the value that reached the optimal DC over the validation set (-0.53 for the raw output predictions).

#### TA-GAN integration in the acquisition loop

The TA-GAN*_Live_* trained for resolution enhancement for live-cell imaging was directly integrated in the imaging acquisition process of the STED microscope (Figure 4a). At the beginning and at the end of each experiments, a FOV of 10 x 10 µm was selected and reference STED and confocal images were acquired. The reference images were used to monitor the dendritic F-actin activity-dependent remodeling in living neurons(Extended data fig. 4). Similarity measurements between the synthetic and real STED images do not show time-dependent changes in the generation accuracy over all imaging sequences (Supplementary Figure 24).

For each time point, as a first step (Table 4), a confocal image of the ROI is acquired to serve as the input to the TA-GAN*_Live_* for the generation of 10 synthetic resolution enhanced images of the ROI (step 2, Table 4). The 10 synthetic images of steps 2 and 5 are generated using different random dropout masks created with the default dropout rate of 0.5 from Srivastava *et al.* [57] and confirmed to be appropriate when applied on GANs by Wieluch and Schwenker [49].

**Table 4:** Steps performed at each time point for automated TA-GAN assistance.

|  |  |
| --- | --- |
| 1 | A confocal image of the FOV (10x10 $\mu\text{m}$) is acquired |
| 2 | Using dropout, 10 synthetic images of the FOV are generated |
| 3 | A sub-region (2x2 $\mu\text{m}$) of highest variability outside the ROI is identified with optical flow |
| 4 | A real STED image of this sub-region is acquired |
| 5 | The confocal image of the FOV and of the STED sub-region are used by the TA-GAN to produce 10 new synthetic STED images of the FOV |
| 6 | The fibers in the ROI (central region of 6 x 6 $\mu\text{m}$) are segmented by U-Net<sub>Live</sub> |
| 7 | The segmentation predictions are used to decide if a STED image should be acquired based on either 1) the mean DC with the segmentation of the last acquired STED, or 2) the variability between the 10 segmentation predictions. |

The third step is the selection of a STED sub-region outside the ROI (step 3, Table 4) which is given as input, along with the confocal FOV, to the TAGAN*_Live_* to account for signal variation in live-cell imaging. In the fourth step, the STED sub-region is acquired on the microscope. Finally, this sub-region (step 5, Table 4) is given as input to the TA-GAN together with the confocal image as described in the previous section. The STED images generated by TA-GAN*_Live_* more closely match the ground truth STED ROI when a STED sub-region is given as input along with the confocal FOV (Supplementary Figure 25).

In our imaging-assistance framework, we choose for step 3 (Table 4) to compute the pair-wise optical flow (OF) between the 10 synthetic images generated with the TA-GAN*_Live_* using dropout. The OF is computed using a Python implementation of the Horn–Schunck method [58] with the Python multiprocessing library, parallelizing the computations on 8 CPUs to increase the computation speed and avoid delays. The OF is computed between each pair of the 10 synthetic images (1-2, 2-3, …). To translate the pixel-wise OF to a region-wise maps, the 500 x 500 pixels OF image was downsampled to a 5 x 5 map using the mean of each 100 x 100 pixels region. The sub-region with the highest mean displacement is imaged with the STED modality. We decided to use OF as a measure of disparity between the synthetic generations, but other measures (e.g. standard deviation, SSIM, mean intensity) could be used for experiments where computation time needs to be minimized (Supplementary Table 2). The sequence of acquiring the full confocal, the STED sub-region, generating 10 synthetic STED images, computing the OF, and taking the decision required around 14 seconds per 500*×*500 pixels regions (10*×*10 *µ* m). Steps 2, 3, 5 and 6 are computed with a graphical processing unit (GPU) to avoid computation induced delays. To do so, the commands from steps 2, 3, 5 and 6 are sent from the microscope’s control computer to a GPU-equipped computer using the Flask [59] web framework Python module. All automated acquisitions use the SpecPy Python library to interface with the Imspector software (Abberior Instruments, Germany).

#### Live-cell imaging decision guidance using the TA-GAN

The TA-GAN*_Live_* predictions are used for decision guidance on the optimal STED and confocal acquisition sequence and applied to the imaging of F-actin remodelling dynamics in cultured hippocampal neurons.

#### TA-GAN assisted monitoring of expected structural change

The proof-of-concept experiment targets the *expected* activity-dependent remodelling of dendritic F-actin rings into fibers [14]. Based on previous findings, the area of F-actin fibers was expected to increase following a neuronal stimulation[14]. The structural remodelling is monitored by comparing the area of segmented F-actin fibers on the synthetic and the reference real STED images. F-actin fibers are segmented on the synthetic STED images by U- Net*_Live_*. At each time point, steps 1-5 are performed as described in Table 4. To decide, following step 5, whether or not a STED image of the full ROI should be acquired, 10 synthetic images of the ROI (acquired with the confocal modality) are generated and segmented by the U-Net*_Live_*. The mean of the 10 segmentation maps is compared to the segmentation map predicted for the last acquired real STED image (reference STED) using the DC metric. A low DC is indicative of changes in the F-actin nanostructures in respect to the reference STED. A full real STED image is acquired if the DC falls below a pre-established threshold of 0.5. The value of 0.5 was chosen by performing several trials on live-cell F-actin imaging. The value of the DC threshold should be adapted to the type of structural remodelling observed. Each time the acquisition of a STED on the full ROI is triggered, the STED reference image is updated for subsequent comparison of the segmentation maps.

#### Monitoring the TA-GAN*_Live_* generator’s variability

The pixel-wise generator’s variability can also be monitored to trigger the imaging of a full ROI with the STED modality. At each time point, steps 1- 5 are performed as described in Table 4. The 10 synthetic images generated at step 5 are segmented by the U-Net*_Live_*, resulting in 10 segmentation maps for F-actin rings and fibers. The 10 segmentation maps of F-actin fibers are binarized and summed. Pixels in the summed segmentation prediction has a value between 0 and 10 (0 when the presence of fibers was predicted in none of the synthetic images and 10 when it is predicted in all). The variability of the generator on the segmentation prediction is evaluated from the summed segmentation prediction. Low variability pixels are the pixels having the same value for at least 80% of the predicted segmentation maps (values of 1-2 (no fibers), 9-10 (fibers) positive counts). High variability pixels are those having positive counts in between (3-8, inclusive). The distribution of high and low variability pixels from the foreground (Figure 4b) is compared for each image. Pixels having 0 positive counts (mostly background) are not considered. The proportion of low variability pixels in the foreground is defined as the variability score (VS). A VS below 0.5 corresponds to images for which the predictions of the U-Net*_Live_* on the 10 synthetic images are consistent for the majority of foreground pixels. If the VS is above 0.5, the 10 synthetic STED images are not consistent and a STED acquisition is triggered. The threshold of 0.5 was chosen because it corresponds to the tipping point where the number of high variability pixels exceeds the number of low variability pixels.

---

**Extended data fig. 1:**
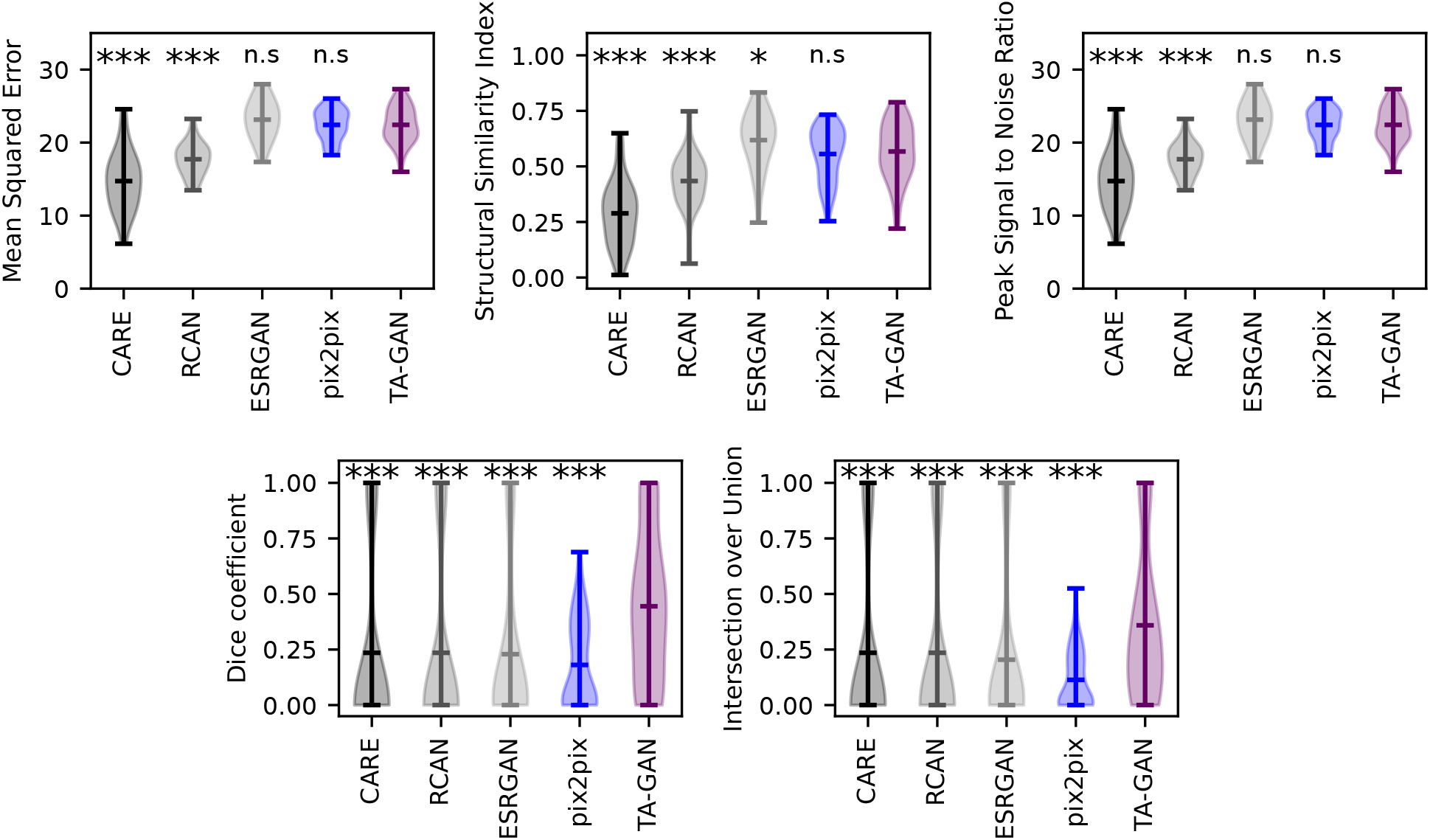
Comparison of TA-GAN*_Ax._* with the resolution enhancement baselines using three image evaluation metrics : 1) Mean squared error (MSE), 2) Structural Similarity Index Measure (SSIM), 3) Peak Signal to Noise Ratio (PSNR), and two segmentation evaluation metrics : 1) Dice Coefficient (DC), 2) Intersection over Union (IOU). For the image metrics, images are normalized to 0-1 using min-max normalization. The segmentation predictions are computed with the U- Net*_Fixed−ax._* on the synthetic images generated with each approach. Metrics are computed using the real STED image and its segmentation by U-Net*_Fixed−ax._* as the reference. The score for DC and IOU is 1 if both the reference and prediction are empty. The performance of the TA-GAN is significantly better than all baseline for both segmentation metrics. For the image similarity metrics, TA-GAN performs significantly better than CARE and RCAN, and is similar to ESRGAN and pix2pix. Statistical analysis : Mann-Whitney U test [50] for the null hypothesis that the distribution underlying the results for each baseline is the same as the distribution underlying the TA-GAN results. (∗∗∗ *p <* 0.001, n.s. *p >* 0.05)

**Extended data fig. 2:**
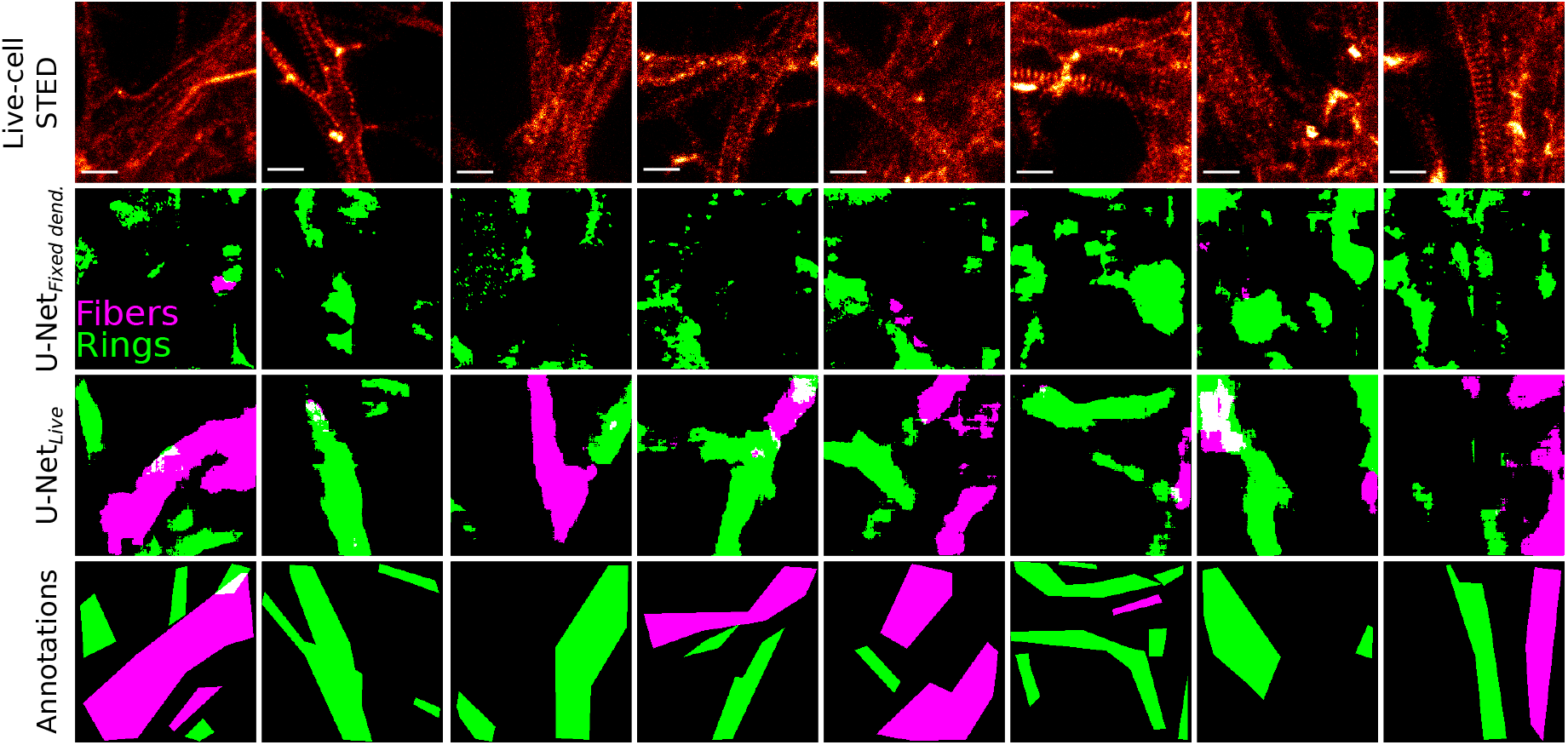
Segmentation predictions by U-Net*_Fixeddend._*[14] and U-Net*_Live_* on randomly sampled images from the *Live F-actin dataset* test set. Annotations were created for visualization purposes and were not used for training U-Net*_Live_*. Only the U-Net*_Live_* trained only on synthetic images from the *Translated F-actin dataset* succeeds in segmenting F-actin nanostructures on real STED images. Scale bars: 1 µm.

**Extended data fig. 3:**
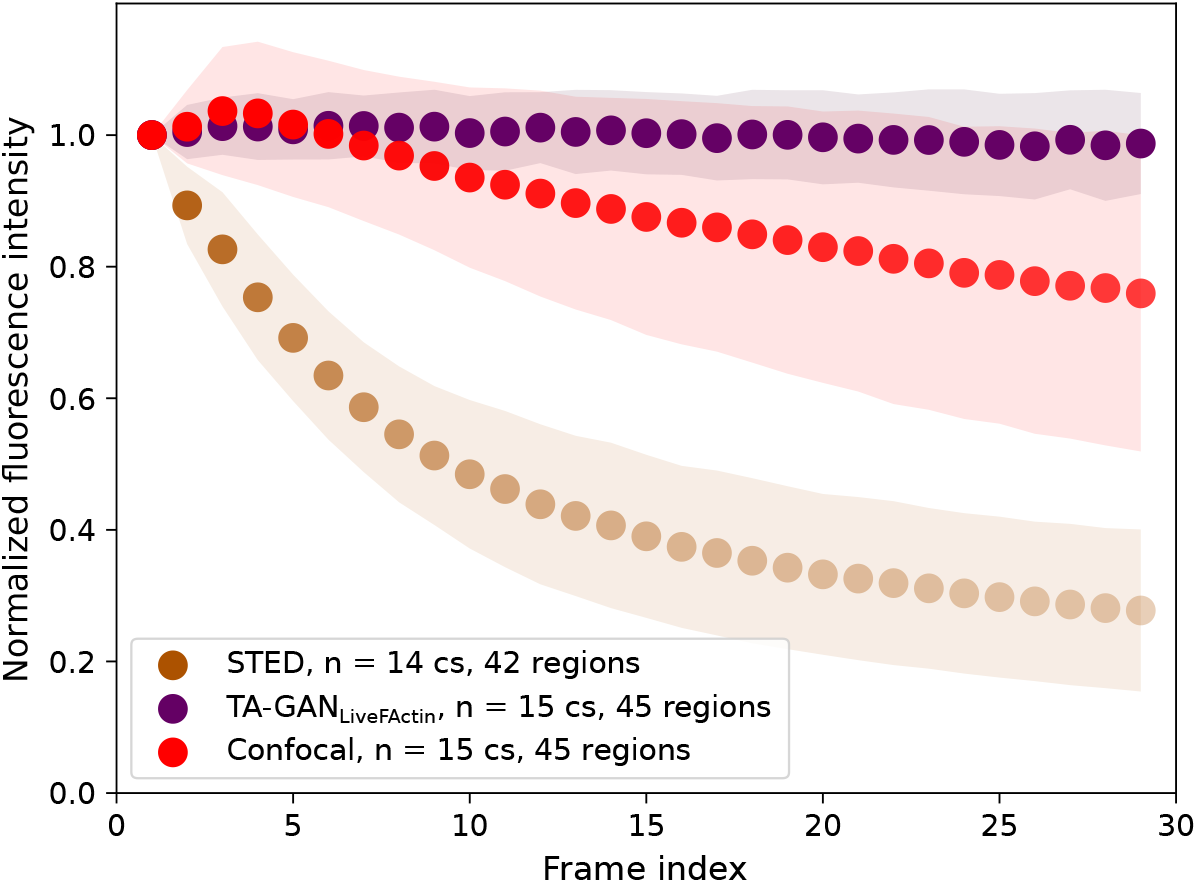
Normalized fluorescence intensity after 30 confocal acquisitions and (red, N=45 regions), associated synthetic STED signal (purple, N=45 regions) over the full FOV in comparison to acquisitions using the STED modality at each frame (orange, N=42 regions). The TA-GAN*_Live_* predictions compensate for the fluorescence intensity decrease in the synthetic STED images.

**Extended data fig. 4:**
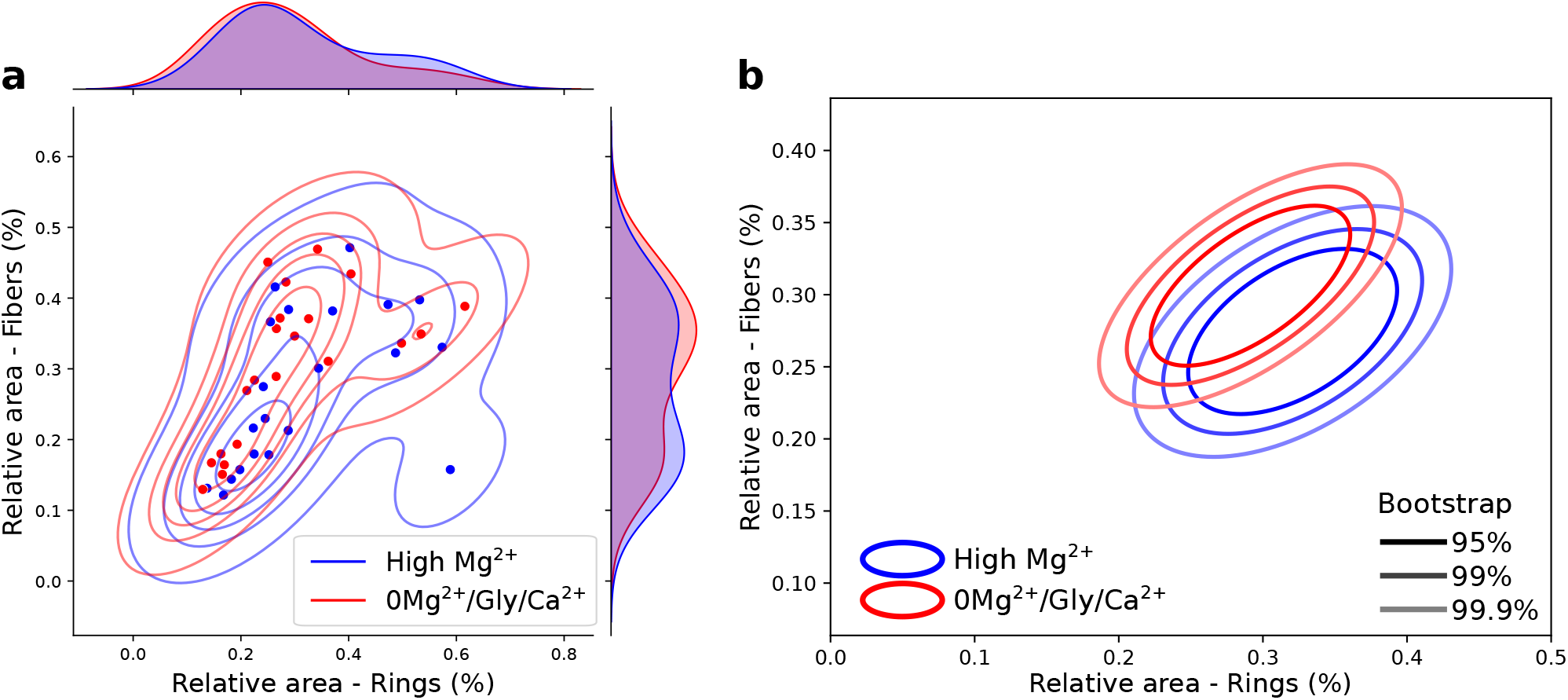
**a,** Kernel density estimate of the F-actin fibers and rings dendritic area distribution for after 30 minutes in a solution reducing neuronal activity (high Mg^2+^/low Ca^2+^, blue) or following a stimulation (0Mg^2+^/Glu/Ca^2+^, from *t* =1-15min, red). **b,** Bootstrapped distributions of the results shown in **a,**. Shown are the regions comprising 95%, 99% and 99.9% of the data point distribution. Following the 0Mg^2+^/Glu/Ca^2+^ stimulation, we observe a small increase in the proportion of F-actin fibers and a decrease in the proportion of rings. High Mg^2+^ N=21, 0Mg^2+^/Glu/Ca^2+^ N=21.

**Extended data fig. 5:**
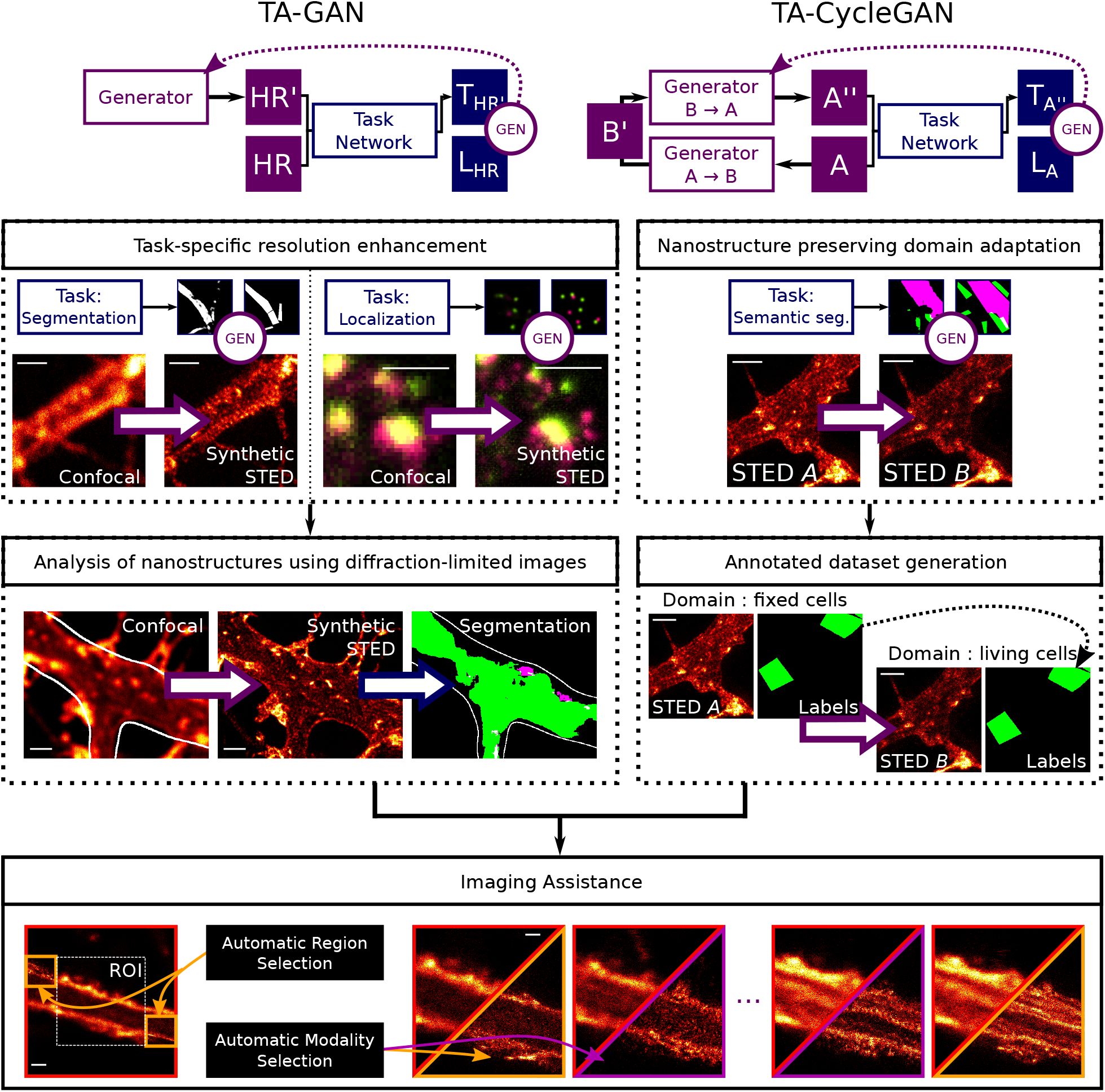
Graphical Abstract. The proposed model has two general use cases: TA-GAN, for paired datasets, and TA-CycleGAN, for unpaired datasets. *Top-left:* The TA-GAN uses a task adapted to each dataset for accurate resolution enhancement. The generation loss (GEN circle) is computed from the comparison between the output of the task network for the synthetic high-resolution image (*T_HR_I* ) and the labels obtained from the ground truth image (*L_HR_*). The loss is backpropagated to the generator (dashed arrow). *Middle-left:* The generated synthetic STED images are used to analyze the distribution of nanostructures that were not resolved in the original confocal image. *Top-right:* Domain adaptation using the TA-CycleGAN enables the generation of large annotated synthetic image datasets from a new domain, even if labels are only available in one domain. The generation loss (GEN circle) is computed from the comparison between the output of the task network for the image and the synthetic version (*T_A_II* ) and the labels obtained from the input domain A image (*L_A_*). The loss is backpropagated to the generator (dashed arrow).*Middle-right:* Labeled datasets from domain A (e.g fixed cells) are adapted to the unlabeled domain B (e.g live cells) to obtain a labeled dataset from domain B, which can be used to train a super-resolution TA-GAN. *Bottom:* Both models can be used for microscopy acquisition guidance. The TA-GAN model, trained using a TA-CycleGAN generated dataset, can automatically identify regions and frames of interest from the low-resolution images. Automatic switching between low- and high-resolution imaging modalities is guided by the TA-GAN*_Live_* predictions. Scale bars: 1 µm.

