## Supplementary Figures for "Resolution Enhancement with a Task-Assisted GAN to Guide Optical Nanoscopy Image Analysis and Acquisition"

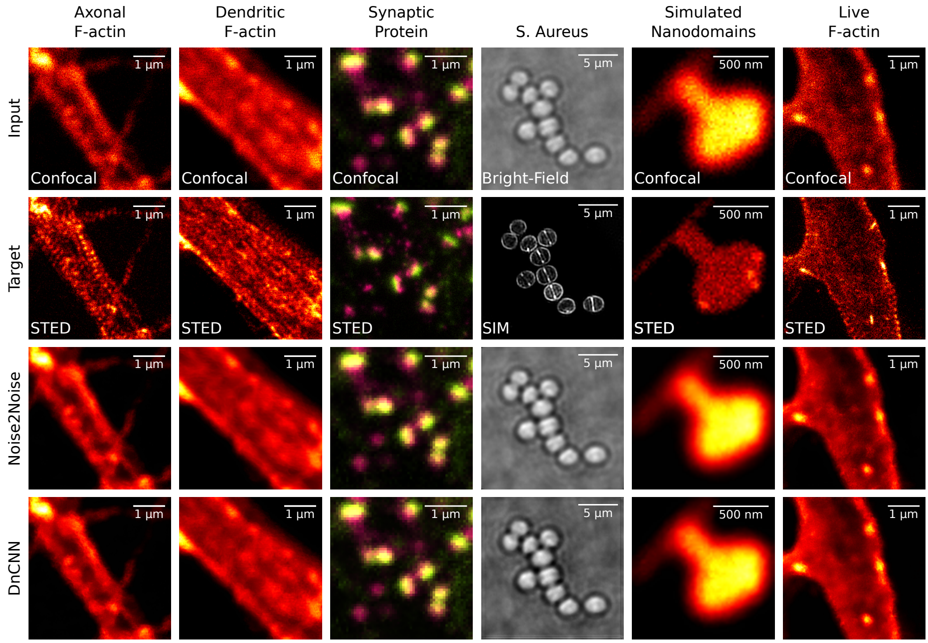


**Supplementary Figure 1:** The denoising approaches Noise2Noise[38] and Denoising CNN[36] cannot generate sub-diffraction structures of interest that are well resolved in the target modality (STED or SIM). Both models are directly applied using the trained weights from [37] since the training of those denoising approaches requires paired images with variable noise levels which were not available with our datasets.

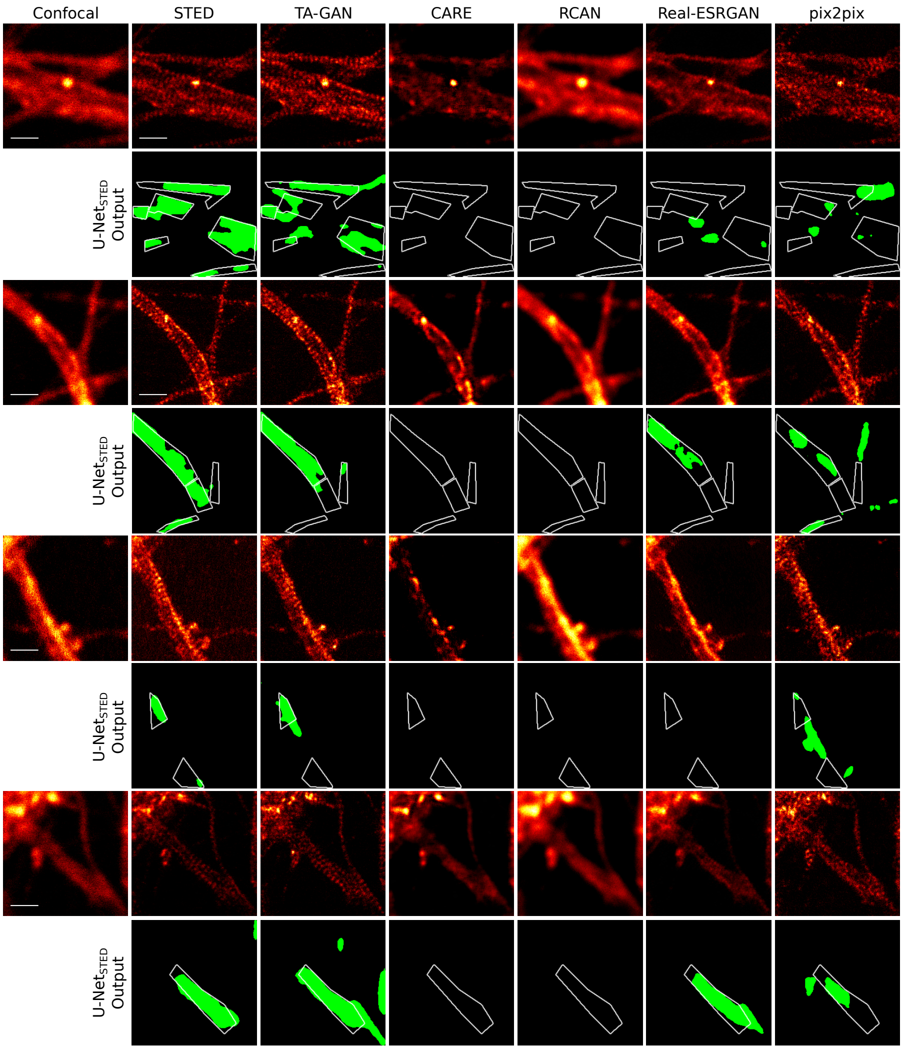


**Supplementary Figure 2:** Comparison of the TA-GAN with other resolution enhancement baselines on randomly selected example from the test set of the *Axonal F-actin dataset*. On all images from the test set, CARE and RCAN fail at generating axonal rings that are recognized by U-Net$_{Fixed-ax.}$. Real-ESRGAN and pix2pix generate periodic F-actin nanostructures that are segmented by U-Net$_{Fixed-ax.}$ but are characterized by a significantly lower Dice Coefficient than TA-GAN$_{Ax.}$ when compared with the segmentation obtained on the corresponding real STED image (Supplementary Figure Extended data fig. 1). Scale bar: 1 µm. The intensity scaling of each image was normalized for display purpose only by its own intensity maximum.

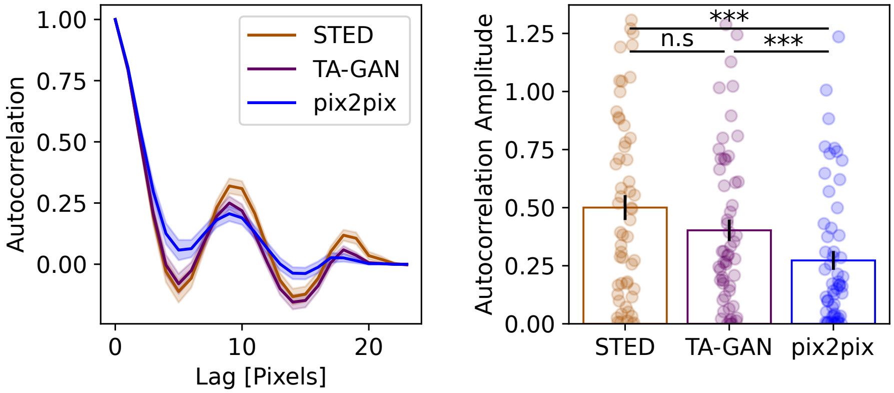


**Supplementary Figure 3:** Left, autocorrelation curve[32] (mean ± SEM) of axonal F-actin in real STED images, TA-GAN$_{Ax.}$ synthetic images, and pix2pix synthetic images. Right, measurement of the corresponding autocorrelation amplitude (mean ± SEM, n = 55 tracings from 26 test images from the *Axonal F-actin dataset*). Statistical analysis: Mann-Whitney U-test (distributions are not drawn from a normal distribution based on the Shapiro-Wilk test for normality). (∗∗∗ $p < 0.001$, n.s. $p > 0.05$)

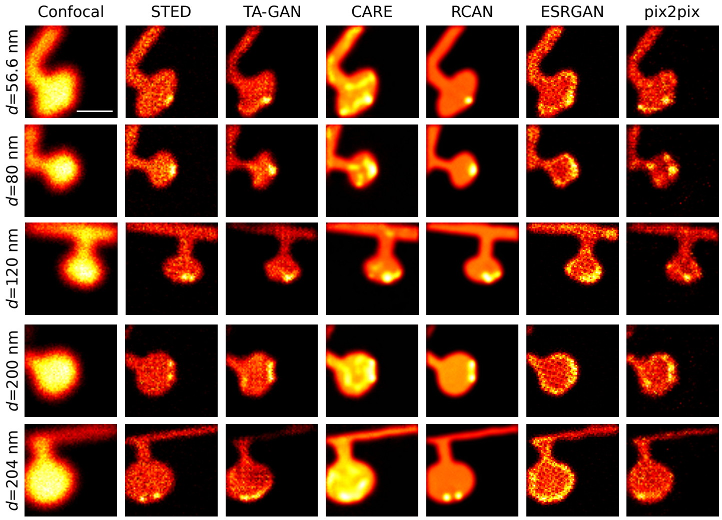


**Supplementary Figure 4:** Examples of synthetic images generated from the two-nanodomains test set of the *Simulated nanoomain dataset*. From top to bottom, the distance $d$ between the pair of nanodomains in the ground truth datamap increases. Scale bar: 500 nm.

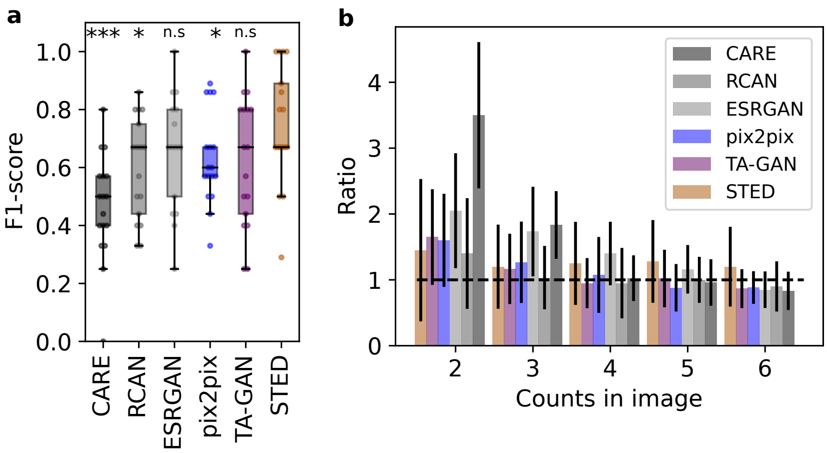


**Supplementary Figure 5: a,** F1-score for the localization of nanodomains on test images with two nanodomains spaced by less than 100 nm. Statistical analysis : Mann-Whitney U test [50] for the null hypothesis that the distributions underlying the results from each method differ from the STED distribution ($*** \ p < 0.001$, $*p < 0.05$, n.s. $p > 0.05$). **b,** Ratio between the predicted and the ground truth number of nanodomains.

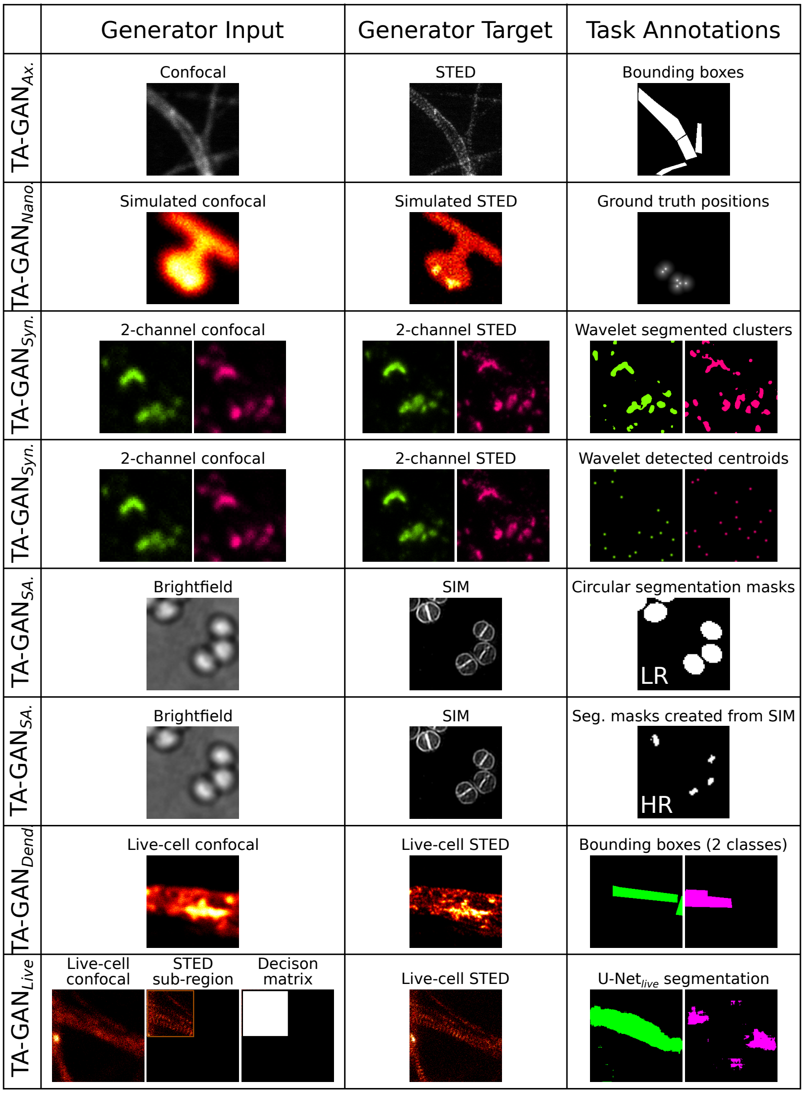


**Supplementary Figure 6:** Inputs and targets for all TA-GAN models.

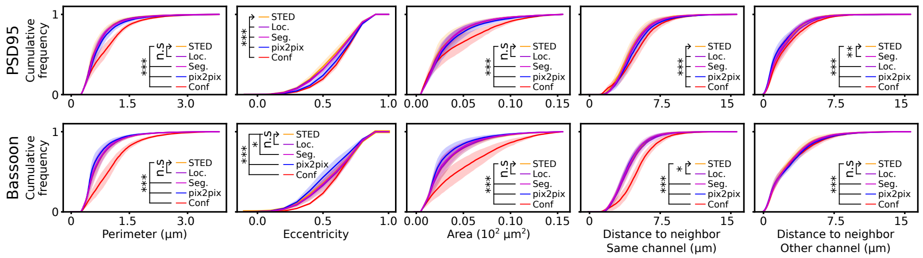


**Supplementary Figure 7:** Cumulative frequency plots of morphological features (perimeter, eccentricity, area, distance to nearest neighbor from the same channel and distance to nearest neighbor from the other channel) of coupled PSD95 and Bassoon clusters [60]. Shown are images obtained with : *red* - confocal, *yellow* - STED, *blue* - pix2pix, *purple* - TA-GAN$_{Syn.}$ using a localization task, and *pink* - TA-GAN$_{Syn.}$ using a segmentation task. Shaded areas show the standard deviation from n = 9 images. The morphological features of the clusters generated by the TA-GAN$_{Syn.}$ trained with the localisation task show the highest similarity with the real STED images. Statistical analysis: two-sample Kolmogorov–Smirnov test for the null hypothesis that the continuous distribution underlying the results for each baseline is the same as the one underlying the STED results ($*** \ p < 0.001$, $** \ p < 0.01$, $*p < 0.05$, n.s. $p > 0.05$)

.

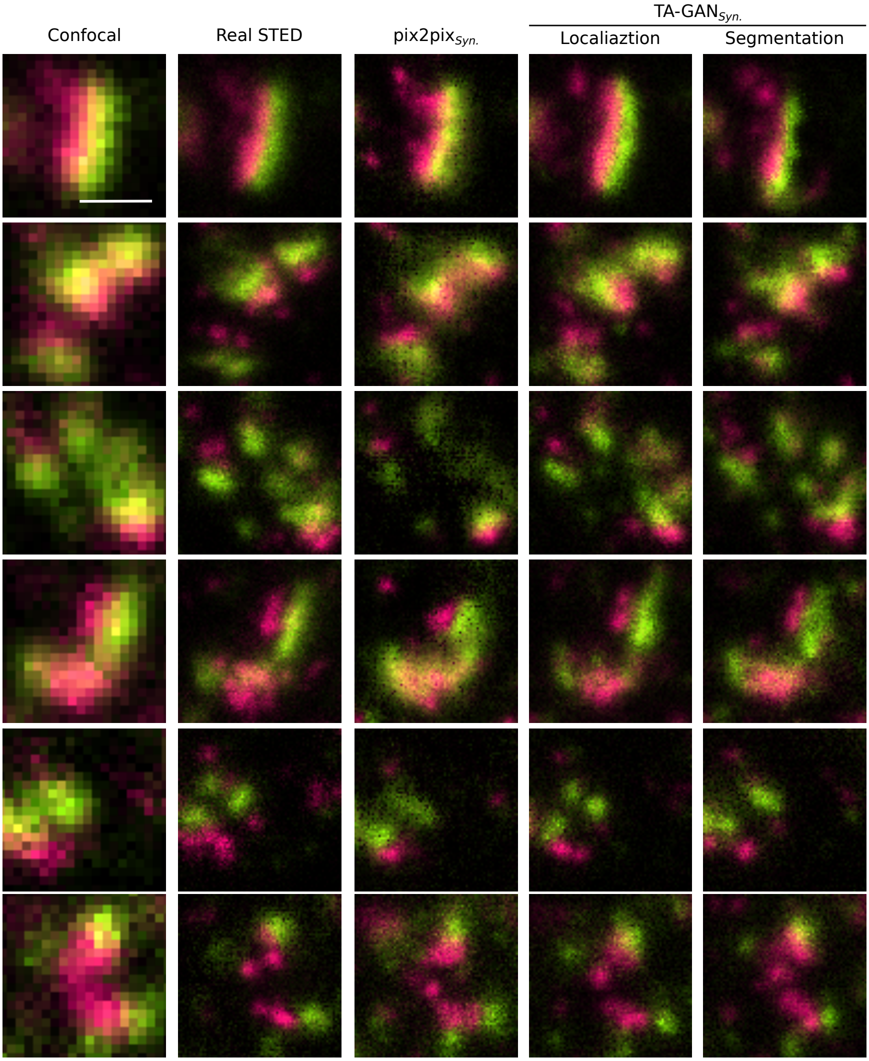


**Supplementary Figure 8: a,** Example of synthetic images generated by the TA-GAN$_{Syn.}$ on the test set of the *Synaptic protein dataset*. Each crop is normalized to its own maximum for better visualization of dim clusters. Scale bar: 500 nm.

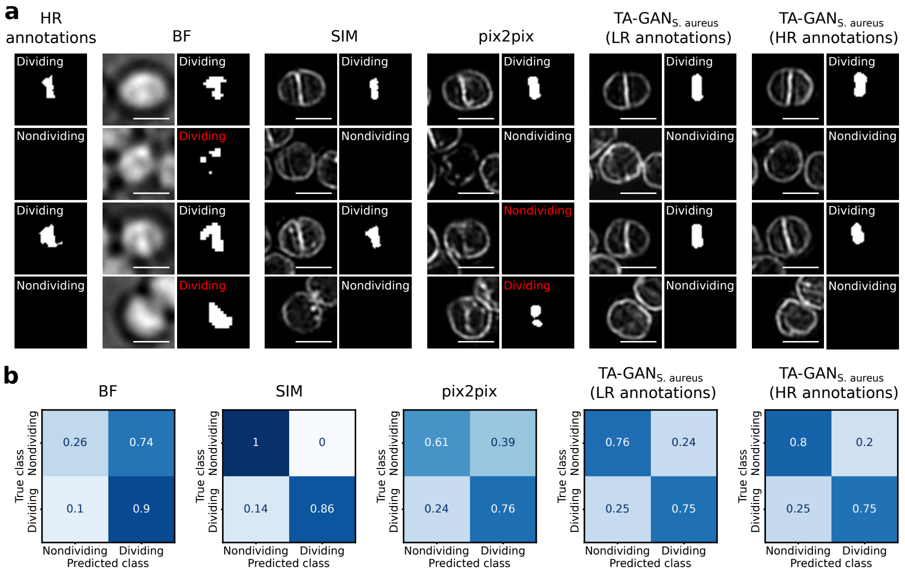


**Supplementary Figure 9:** Classification task developed for the evaluation of TA-GAN$_{SA.}$. **a,** Images and corresponding annotations obtained on the test set. The ground truth annotations are obtained from the high resolution (HR) SIM images. The threshold-based segmentation of the BF images does not allow to reliably identify dividing cell boundaries. The ResNet$_{SA.}$ predicts reliable segmentation masks for dividing cell boundaries on the SIM images. The ResNet$_{SA.}$ is applied for the segmentation of dividing cell boundaries on synthetic SIM images generated with pix2pix$_{SA.}$ and with the TA-GAN$_{SA.}$ trained either with whole-cell low-resolution (LR) annotations from[44] or the ground truth HR annotations shown in the first column. The ResNet$_{SA.}$ trained on real images of the *S. aureus dataset* predicts reliable segmentation masks of the dividing cell boundaries on the synthetic images generated with both training schemes of the TA-GAN$_{SA.}$, but fails at segmenting the structures on the images generated with pix2pix. The class is predicted by applying a threshold on the number of segmented pixels (Methods). All images are normalized to their own maximum. Scale bars: 1 µm. **b,** Confusion matrices for the classification of dividing or non-dividing cells. Classification accuracy : BF 51.0%, SIM 94.4%, pix2pix 66.9%, TA-GAN$_{SA.}$ trained with LR annotations 75.9%, TA-GAN$_{SA.}$ trained with HR annotations 78.3%.

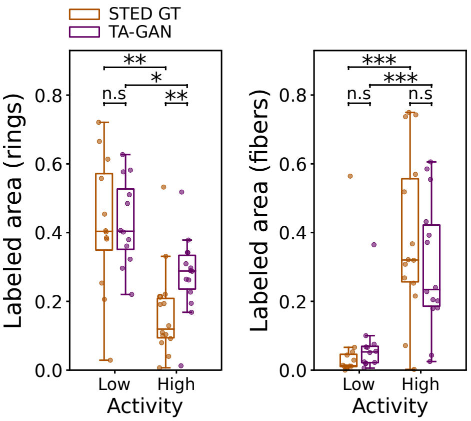


**Supplementary Figure 10:** The proportion of dendritic F-actin rings (left) is significantly larger at low neuronal activity (high $Mg^{2+}$, low $Ca^{2+}$ blocking solution) compared to high activity ($0Mg^{2+}$/Gly/Bicc) in both real (orange) and TA-GAN$_{Dend.}$ generated synthetic (purple) STED images (STED: $p = 0.0007$, TA-GAN$_{Dend.}$: $p = 0.004$). The opposite is observed for F-Actin fibers (right) for both the real ($p = 0.0006$) and the synthetic STED images ($p = 0.0009$). Statistical analysis: Mann-Whitney U test [50] for the null hypothesis that the distributions underlying the compared data are different ($*** \ p < 0.001$, $** \ p < 0.01$, $*p < 0.05$, n.s. $p > 0.05$).

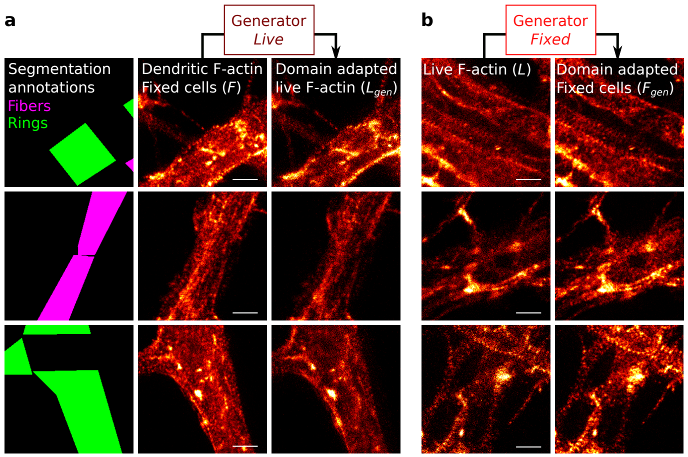


**Supplementary Figure 11:** Fixed-cell to live-cell imaging domain adaptation. **a,** Real STED images of fixed cells are translated to synthetic images from the live-cell imaging domain by $\text{Generator}_{Fixed}$. Following the image translation, the manual segmentation masks associated with the real STED images of fixed cells are associated to the synthetic live-cell images. **b,** Real live-cell images of F-actin are translated to the fixed-cell imaging domain with $\text{Generator}_{Live}$. Scale bars: 1 µm.

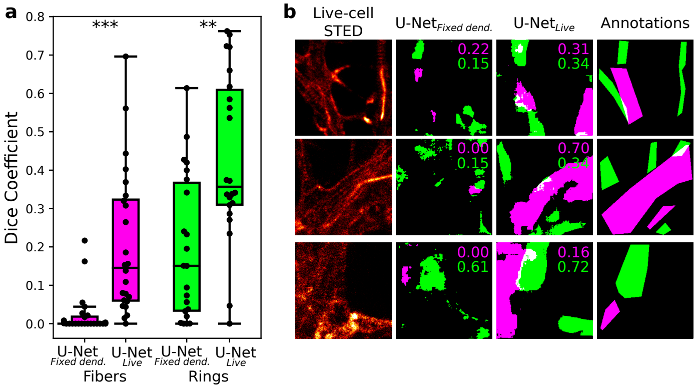


**Supplementary Figure 12:** Measured dice coefficient (DC) for the segmentation of F-actin rings and fibers in real live-cell STED images. **a,** The DC is computed between the predictions of the U-Net segmentation networks and manual polygonal bounding box on live-cell STED images of size (28 images of size 256 x 256 pixels, 5.12 x 5.12 µm.) that were annotated by an expert. 22 images contained F-actin rings and 24 images F-actin fibers. The DC is significantly improved for the segmentation of F-actin nanostructures on live-cell STED images with U-Net$_{Live}$, trained with synthetic images translated to the live-cell domain from the *Translated F-actin dataset*($L'$, mean $DC_{fibers}$ = 0.20, $DC_{rings}$ = 0.43), in comparison with the U-Net$_{Fixed}$, a U-Net trained with the same original fixed-cells STED images from the *Dendritic F-actin dataset* ($F$, mean $DC_{fibers}$ = 0.02, $DC_{rings}$ = 0.20). Statistical analysis: two-sample Kolmogorov–Smirnov test ($*** \; p < 0.001$, $** \; p < 0.01$). **b,** Examples of predicted segmentation masks from the *Live F-actin dataset* test set with their individual DC written in the top right corner for F-actin fibers (magenta) and rings (green). From left to right: live-cell STED image, predicted segmentation from the U-Net$_{Fixed}$ trained on fixed-cell images ($F$), predicted segmentation from U-Net$_{Live}$ ($L'$, and manual expert annotations. Top row: best DC on fibers obtained with U-Net$_{Fixed}$. Middle row: best DC on fibers obtained with U-Net$_{Live}$. Bottom row: best DC on rings obtained with U-Net$_{Fixed}$.

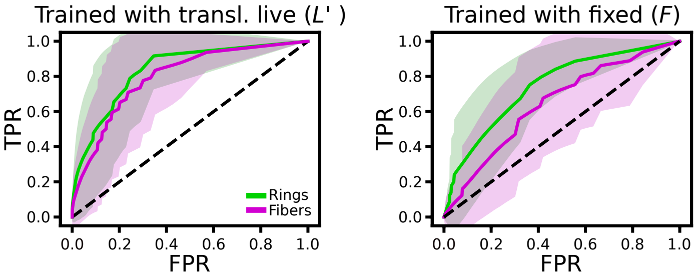


**Supplementary Figure 13:** ROC curves computed over the test set (N=28) of synthetic images from the live-cell domain from the live-cell domain, using the manual annotations as ground truth. The TA-CycleGAN translates real fixed-cell STED images from the *Dendritic F-actin dataset* into corresponding synthetic images from the live-cell STED imaging domain (*Translated F-actin dataset*). Left: the U-Net$_{Live}$ is trained on the *Translated F-actin dataset* ($L'$). On the right: U-Net$_{Fixed}$ is trained on the *Dendritic F-actin dataset* ($F$). The results are reported for F-actin fibers (magenta) and rings (green). The shaded region corresponds to the standard deviation. Dashed line represents the worst achievable performance (random classifier, AUROC=0.5).

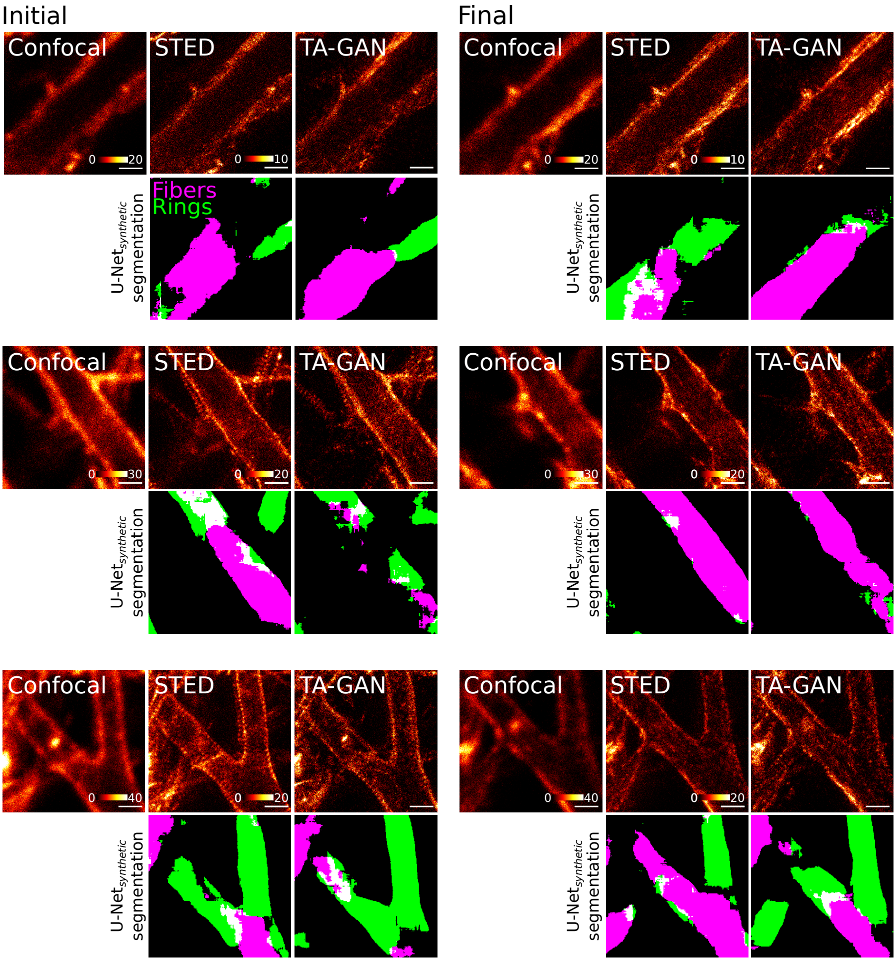


**Supplementary Figure 14:** Live-cell imaging sequences using the TA-GAN$_{Live}$ before (left, *initial*) and after (right, *final*) neuronal stimulation.

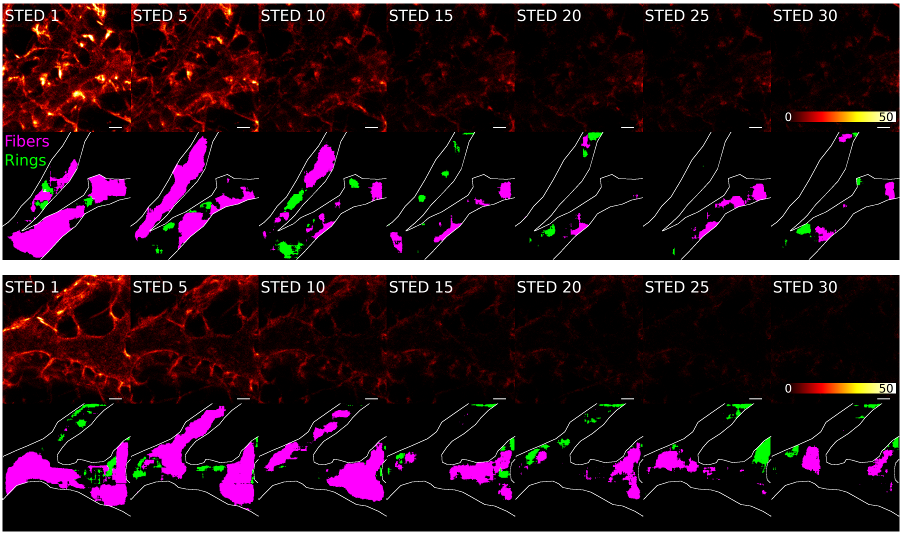


**Supplementary Figure 15:** Two sequences of 30 STED acquisitions are used to evaluate the photobleaching. The STED images at frames 1, 5, 10, 15, 20, 25, and 30 are displayed using the same intensity scale. The segmentation prediction is obtained with the U-Net$_{Live}$. The signal in the STED images quickly degrades after a few acquisitions, and U-Net$_{Live}$ fails at accurately identifying the F-actin nanostructures. Scale bars: 1 µm.

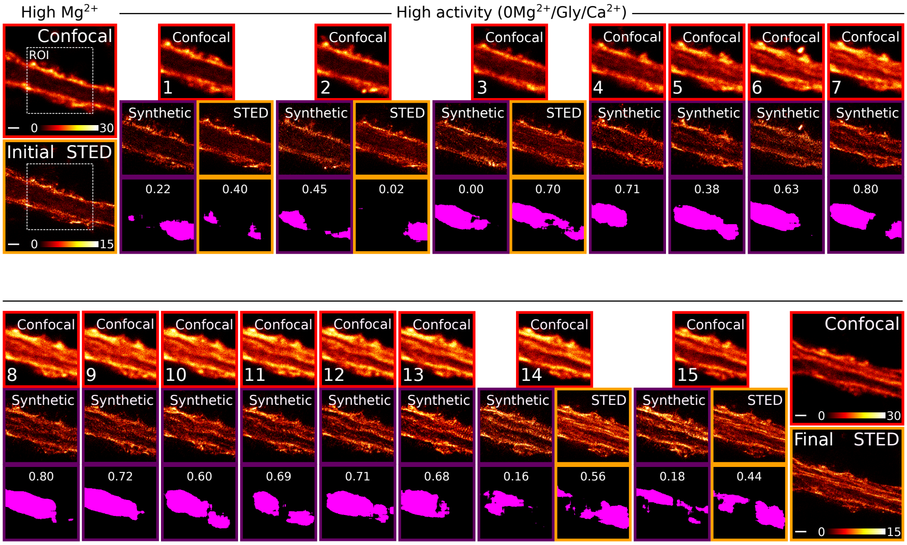


**Supplementary Figure 16:** Images acquired at all time points for the time series shown in Figure 4**a**. First row shows the confocal images (red), second row shows the STED images which are either synthetic (purple) or real (orange), third row shows the segmentation of fibers by applying the U-Net$_{Live}$ on the STED (synthetic or real) images. The DC computed with the segmentation of the most recent real reference STED as the ground truth is displayed on each segmentation map. The intensity scale is constant for all confocal images (0-30 counts) and for all STED images (0-15 counts). Scale bars: 1 µm.

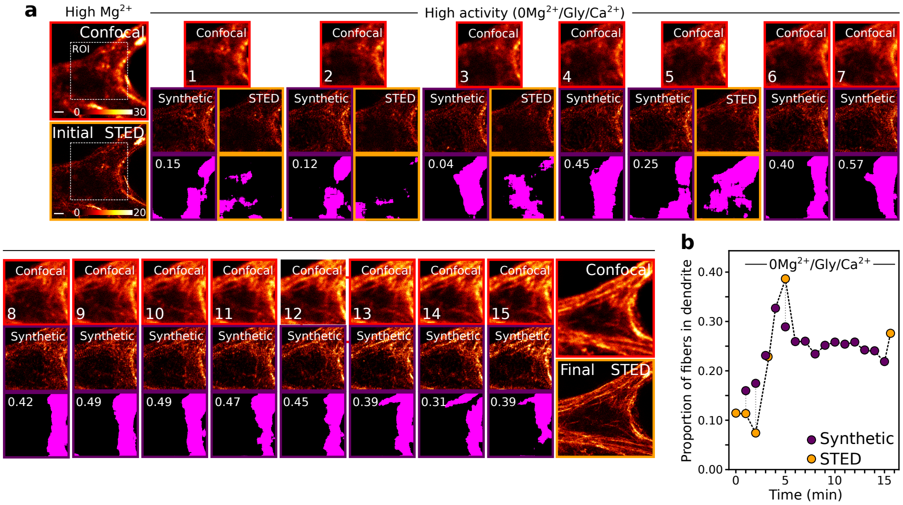


**Supplementary Figure 17:** Additional example showing the prediction of F-actin remodelling with TA-GAN$_{Live}$. **a,** Live-cell imaging of dendritic F-actin before (initial), during (frames 1-15) and after (final) application of a 0Mg$^{2+}$/Gly/Ca$^{2+}$ solution, which promotes synaptic NMDA receptor activity. Shown are the confocal images (red, top row), synthetic STED images (purple, middle row), and real STED images when acquired (orange, middle row), and corresponding segmentation masks for F-actin fibers (bottom row). **b,** The U-Net$_{Live}$ is used to compute the proportion of dendritic F-actin fibers on either the real STED (orange) or the synthetic STED (purple). In this specific case, STED acquisitions are triggered after 1, 2, 3, and 5 minutes, allowing to image in nanoscale resolution the increase in the proportion of F-actin fibers. The proportion of F-actin fibers remains stable for the following time points (6-15) and no STED images are triggered. Scale bars: 1 µm.

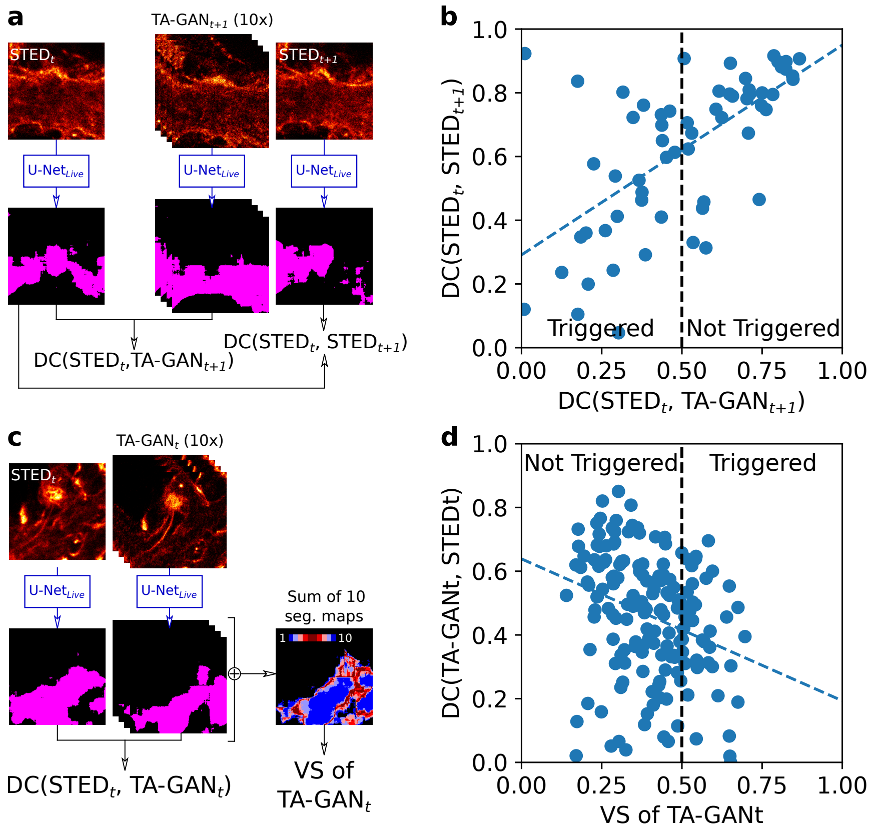


**Supplementary Figure 18: a,** The Dice coefficient (DC) between the real STED at time $t$ and the TA-GAN synthetic STED at time $t$+1 (one minute later) is computed. The DC between the real STED from time $t$ and the next STED from time $t+1$ is computed. **b,** Both DC computed in **a** are compared. The change in the segmentation between the two real STED images at times $t$ and $t$+1 (y-axis) is positively correlated with the similarity between the real STED image at time $t$ and the synthetic STED image at time $t+1$ (x-axis, $r^2 = 0.397$, $p \sim 10^{-8}$). The correlation implies that the DC threshold can be adjusted to the objective of a given experiment: for example increasing the threshold would trigger less acquisition of STED images and only at frames where the amount of change is higher. **c,** The DC between the real and synthetic STED images at time $t$ is computed. The segmentation maps of the 10 synthetic STED images are used to compute the variability score (VS) at time $t$. **d,** The DC and the VS computed in **c,** are compared. The changes between the segmentation of the real and synthetic STED images at time $t$ (y-axis) is negatively correlated with the variability score (x-axis, $r^2 = 0.156$, $p = 0.0018$). The correlation implies that the VS is a good measure of the uncertainty of the TA-GAN, since higher VS correlates with higher disparity with the real STED segmentation. Statistical analysis : Wald Test with t-distribution of the test statistic for the null hypothesis that the correlation is zero. The analysis was performed on control sequences of two frames with neuronal stimulation (N=60) for which the confocal and STED images were always acquired at time steps $t$ and $t+1$.

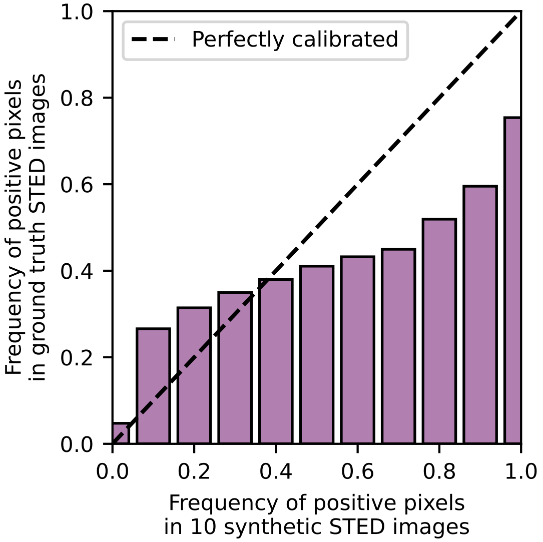


**Supplementary Figure 19:** Reliability diagram for the segmentation of F-actin fibers on the *Translated F-actin dataset*. Positive pixels are pixels identified as fibers by U-Net$_{Live}$. Predicted probability (x-axis) is obtained from the segmentation of 10 synthetic STED images generated by TA-GAN$_{Live}$. Observed probability (y-axis) is obtained from the segmentation of the corresponding ground truth STED images. N=162 ROIs of size 300 x 300 pixels. The fact that the function is monotonic validates the use of this metric to trigger a STED acquisition.

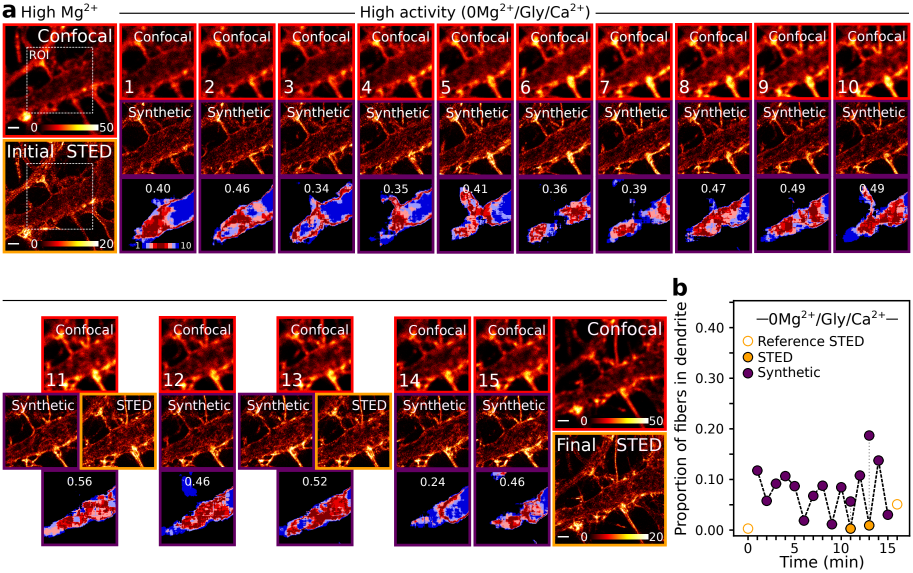


**Supplementary Figure 20: a,** Images at all time points for the time series shown in Figure 4**f**. First row shows the confocal images (red), second row shows the STED images which are either synthetic (purple) or real (orange), third row shows the sum of 10 segmentation maps of F-actin fibers obtained by applying the U-Net$_{Live}$ the STED images. Blue pixels have low variability and red pixels have high variability. The the variability scores are displayed on each segmentation maps. The intensity scale is constant for all confocal images (0-50 counts) and for all STED images (0-20 counts). Scale bars: 1 µm. **b,** Proportion of fibers on the dendrite on either the real STED (orange) or the synthetic STED (purple) images. Initial and final reference STED images (empty orange bullets) are acquired at each round. In this specific case, there is no activity-dependent increase.

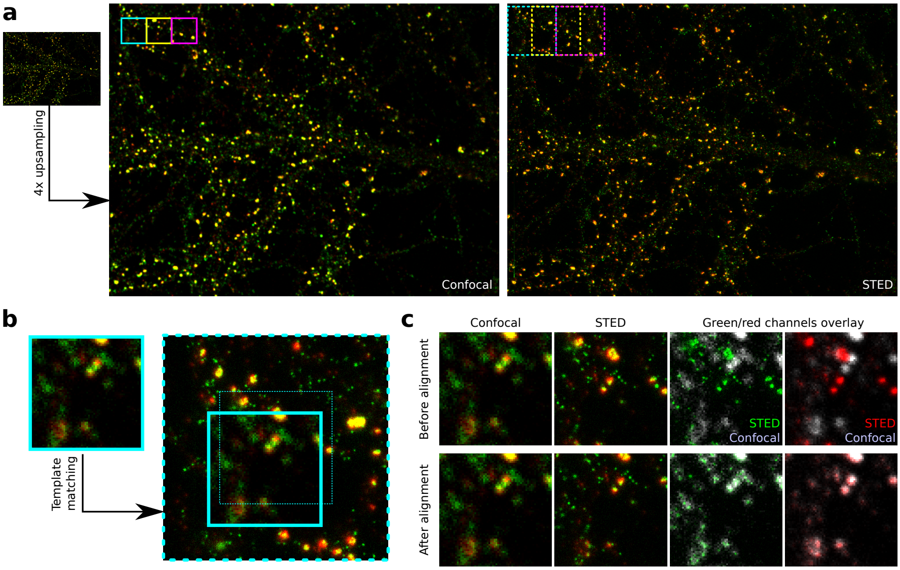


**Supplementary Figure 21:** Registration method to align confocal and STED image pairs from the *Synaptic protein dataset.* **a,** The confocal image is upsampled with nearest-neighbor interpolation to match the size of the STED image. The three regions identified (green, yellow and magenta squares) show the sliding window method used to iteratively choose sections to match. **b,** For each 512 x 512 px square region of the STED image, the centered 256 x 256 px square region from the confocal image is matched using the *OpenCV* [61] template matching library. The dashed square identifies the center of the region, and the full line identifies the region which matches the confocal crop. **c**) The confocal crop, the STED corresponding crop, and a channel-wise overlay of the two modalities before (top) and after (bottom) alignment.

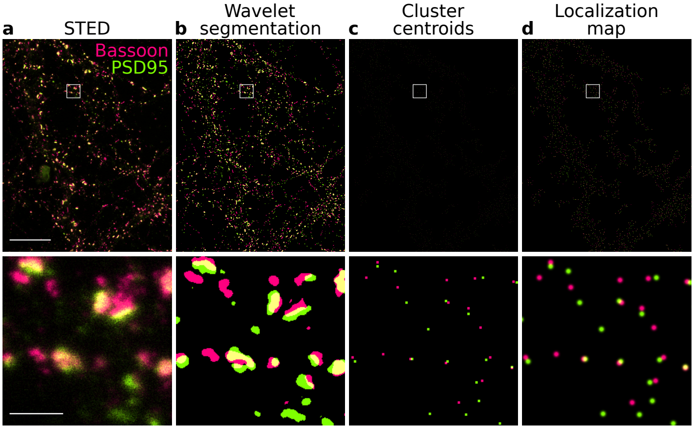


**Supplementary Figure 22:** Localization maps of synaptic protein clusters. **a,** Two-color STED image of PSD95-Alexa594 (green) and Bassoon-STAR635 (magenta) in fixed neurons [41]. **b,** Wavelet segmentation of Bassoon (pink) and PSD95 (green) clusters. **c,** Weighted centroids obtained from the segmentation of individual clusters. **d,** Final localization maps generated by applying a Gaussian filter on the weighted centroids (Methods). Scale bars: 10 µm (top), 1 µm (bottom).

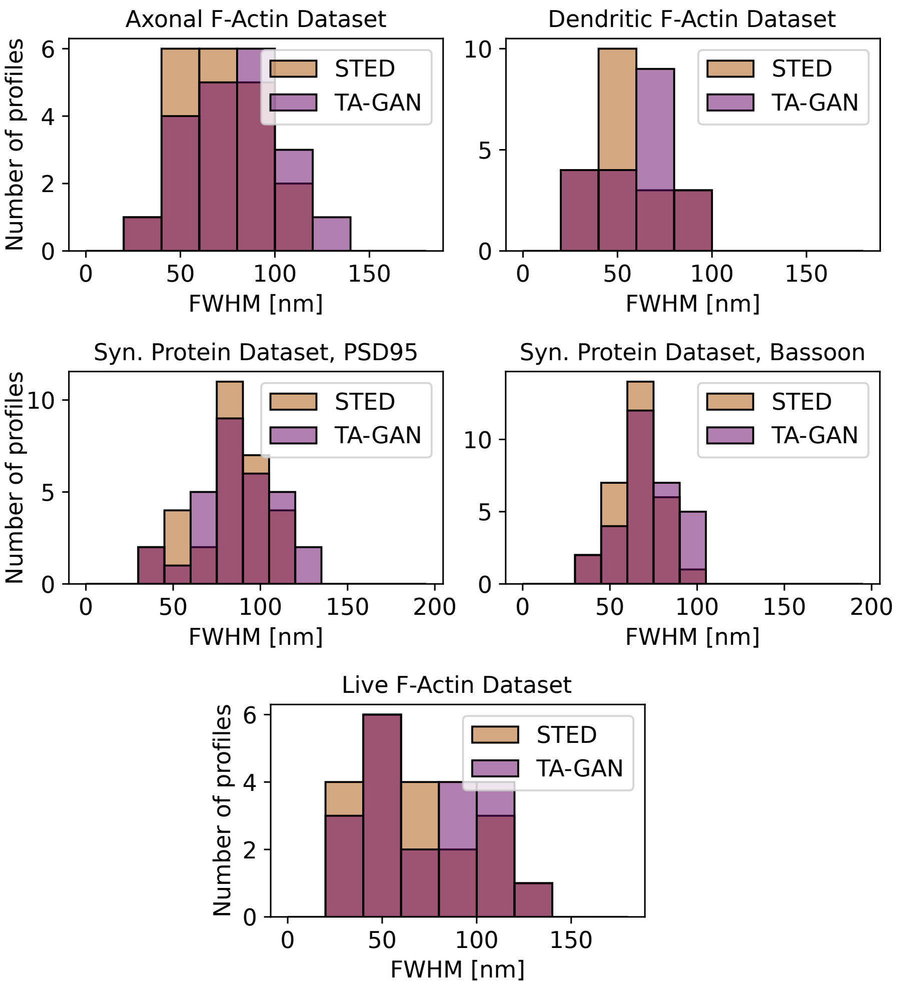


**Supplementary Figure 23:** Distribution of the measured Full Width at Half Maximum (FWHM) on fitted Lorentzian line profiles traced on randomly chosen test images for each dataset (N=30 from 2 images for the *Synaptic protein dataset*, N=20 from 5 images for the *Axonal F-actin dataset*, N=20 from 2 images for the *Dendritic F-actin dataset*). TA-GAN and STED distributions are not statistically different for all datasets (p>0.05). Statistical analysis : Mann-Withney U-test for the *dendritic F-actin dataset* and the *Live F-actin dataset* (distributions are not drawn from a normal distribution based on the Shapiro-Wilk test for normality), independent 2 sample t-test for the *Axonal F-actin dataset* and the *Synaptic Proteins dataset* (equality of variances is verified by the F-test with a significance level of 5%). All bin widths correspond to the pixel size.

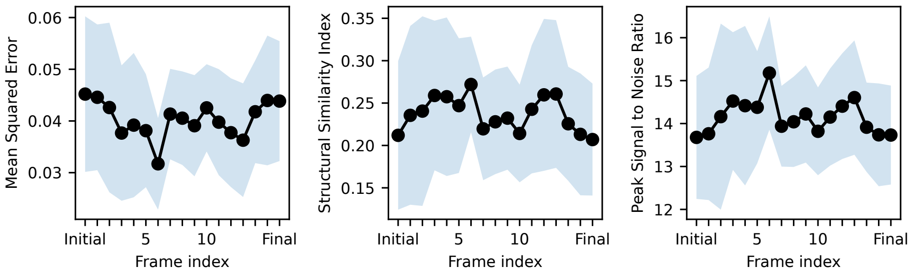


**Supplementary Figure 24:** Comparison between the TA-GAN$_{live}$ synthetic STED images and the corresponding ground truth STED images using the mean squared error, peak signal to noise ratio, and structural similarity). The images (synthetic and STED) are normalized prior to computing the metrics. For the three metrics, the median of the distributions are not statistically different. Solid lines shows the mean value at each frame and the shaded area corresponds to the standard deviation. Statistical analysis : Kruskal-Wallis H-test for independent samples (distributions are not drawn from a normal distribution based on the Shapiro-Wilk test for normality).

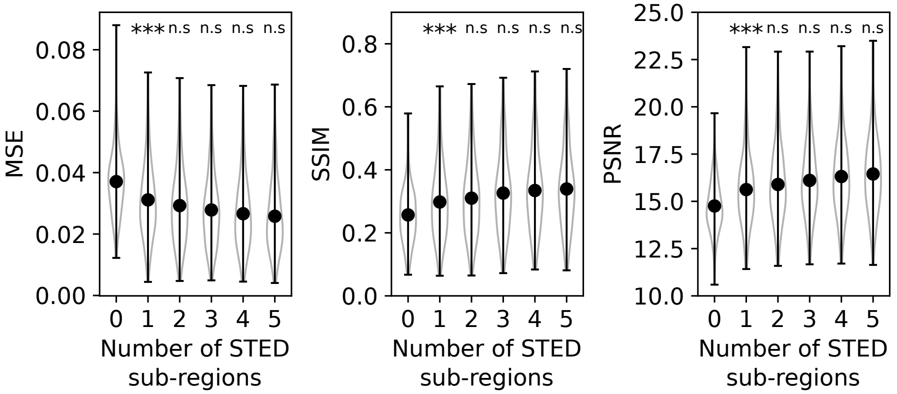


**Supplementary Figure 25:** Generation performance as a function of the number of STED sub-regions given as input to the generator for live-cell imaging assistance. Black dots are the distribution means. Adding one live-cell STED sub-region (100 x 100 pixels) to the input (500 x 500 pixels) of the generator at inference time significantly improves the generation accuracy in terms of MSE, SSIM, and PSNR in the central ROI (300x x 300 pixels). The generation accuracy does not further improve when using more than one STED sub-region. Statistical analysis: Mann-Whitney U-Test for the null hypothesis that the distribution underlying the values for $n$ sub-regions is different to the distribution underlying the values for $n-1$ sub-regions. N=168 images. $***$ $p < 0.001$, $**$ $p < 0.01$, n.s $p > 0.05$.

| | TA-GAN resolution [nm] | STED resolution [nm] |
|---|---|---|
| **Axonal F-actin (far red)** | 75.3 ± 5.4 | 72.8 ± 4.6 |
| **Dendritic F-actin (far red)** | 60.5 ± 4.1 | 57.0 ± 3.5 |
| **Synaptic Proteins** | | |
| Bassoon (Far red) | 72.5 ± 2.8 | 67.3 ± 2.3 |
| PSD95 (red) | 87.2 ± 4.0 | 80.7 ± 3.6 |
| **Live F-actin (far red)** | 73.2 ± 6.3 | 69.3 ± 6.5 |

**Supplementary Table 1:** Resolution (mean ± standard error from the mean) approximated by measuring the Full Width at Half Maximum (FWHM) on fitted Lorentzian line profiles traced on random test images for each dataset (N=30 from 2 images for the *synaptic protein dataset*, N=20 from 5 images for the *Axonal F-actin dataset*, N=20 from 2 images for the *dendritic F-actin dataset*, N=20 from 3 images for the *Live F-actin dataset*). The corresponding resolution measured for confocal images (pixel size 20 nm) is 255.2 ± 1.9 nm for the far red channel (N = 21, 640 nm excitation) and 226.7 ±1.9 nm for the red channel (N = 34, 561 nm excitation).

| Sub-region selection method | **MSE** | **PSNR** | **SSIM** | **Sub-regions used out of 16** |
|---|---|---|---|---|
| **None** | 0.00762 ± 0.00440 | 21.73 ± 2.11 | 0.518 ± 0.123 | N/A |
| **Random** | 0.00696 ± 0.00411 | 22.17 ± 2.22 | 0.535 ± 0.124 | 9.9 ± 1.2 |
| **Min. intensity** | 0.00697 ± 0.00391 | 22.13 ± 2.17 | 0.534 ± 0.123 | 1.9 ± 0.9 |
| **Max. intensity** | 0.00676 ± 0.00398 | 22.16 ± 2.15 | 0.534 ± 0.125 | 2.6 ± 1.1 |
| **Normalized STD** | 0.00692 ± 0.00397 | 22.16 ± 2.15 | 0.536 ± 0.125 | 3.7 ± 1.5 |
| **SSIM** | 0.00676 ± 0.00392 | 22.27 ± 2.20 | 0.544 ± 0.127 | 3.2 ± 1.7 |
| **Optical Flow** | 0.00683 ± 0.00387 | 22.23 ± 2.20 | 0.542 ± 0.126 | 7.3 ± 1.7 |

**Supplementary Table 2:** Generation performance and diversity as a function of the method used to select the STED sub-region given as input to the generator. The metrics evaluated are the averaged Mean Squared Error (MSE), Structural Similarity Index Metric (SSIM), Peak Signal to Noise Ratio (PSNR), and the number of different STED sub-regions selected out of 16 over 15 frames (N=18 sequences). Similarity metrics are computed in the central 300 x 300 px ROI for N=168 pairs of synthetic and real STED images. Achieving a high diversity in the acquired STED sub-regions without compromising generation performance is essential to limit the photobleaching effects that would occur if using the same STED sub-region at each frame.
